# Cross-Dataset Reanalysis of Mouse Alzheimer Disease Single-Cell RNA-Sequencing Data Reveals a Reproducible Innate Immune Activation Signature

**DOI:** 10.64898/2026.09.30.755581

**Authors:** Aryan Anand, Elamathi Natarajan

## Abstract

Single-cell RNA sequencing (scRNA-seq) studies of Alzheimer’s disease (AD) mouse models have proliferated across independent laboratories, brain regions, and platforms, yet the cross-study reproducibility of transcriptional signatures derived from such data remains underexplored. We reanalyzed six publicly available mouse brain scRNA-seq datasets (GSE140399, GSE140510, GSE143758, GSE208683, GSE214244, and GSE296027; 360,620 cells, 102 annotated clusters) using a standardized pipeline (Figure 1) comprising quality control, Leiden clustering, confidence-tiered marker-based cell-type annotation, sample-level pseudobulk differential expression, cross-dataset consistency analysis, and pathway enrichment. Pseudobulk differential expression, performed on 32 samples (16 AD, 16 Control) from five of the six datasets, identified 301 significant genes (FDR-adjusted p < 0.05) out of 20,629 tested, of which 290 (96.3%) were upregulated and 11 (3.7%) downregulated in AD. The upregulated set was dominated by innate immune and interferon-response genes (Ccl3, Ccl4, Ccl6, Itgax, Cst7, Tlr4, Trem2) and showed extremely strong enrichment for immune system and innate immune system pathways (Reactome, p = 1.9 × 10□□³ and 2.1 × 10□²□ respectively), phagosome and osteoclast differentiation (KEGG), and interferon alpha/beta signaling. Of the 301 significant genes, 235 (78%) showed the same direction of effect across all datasets in which they were tested. The sixth dataset, GSE296027, was excluded from differential expression analysis due to a gene-identifier corruption artifact introduced during pseudobulk aggregation (detailed in Methods and Limitations) but contributed valid cell-type composition and microglial subclustering data, independently recovering a disease-associated microglia-like state. These findings indicate that a reproducible, strongly immune/interferon-skewed transcriptional signature, rather than a balanced up/down transcriptional shift, characterizes the mouse AD models analyzed here.

## 1. Introduction

Microglia, the resident innate immune cells of the central nervous system (Prinz et al., 2019), adopt a distinct disease-associated transcriptional state (DAM) in mouse models of Alzheimer’s disease, characterized by downregulation of homeostatic markers such as P2ry12, Tmem119, and Cx3cr1 (Butovsky et al., 2014; Bennett et al., 2016), and upregulation of Trem2, Apoe, and phagocytosis/lipid-metabolism-related genes (Keren-Shaul et al., 2017; Krasemann et al., 2017; Deczkowska et al., 2018; Butovsky and Weiner, 2018). This state has been described using varied experimental designs, mouse models, brain regions, and sequencing technologies (Mathys et al., 2017; Sala Frigerio et al., 2019; Zhou et al., 2020; Chen and Colonna, 2021), raising the question of how reproducible these signatures are when a single analytical framework is applied independently across studies.

Public repositories such as the Gene Expression Omnibus (GEO; Edgar et al., 2002; Barrett et al., 2013) now host dozens of single-cell datasets relevant to AD, representing a substantial but underused resource for cross-study reproducibility analysis. A naive approach, concatenating all available datasets into a single integrated object, is difficult to defend given the heterogeneity of sample sizes, cell numbers, tissues, ages, and experimental designs across studies. We instead adopt a federated reanalysis strategy: each dataset is processed independently through a standardized pipeline, and reproducibility is assessed post hoc by comparing the direction and significance of gene-level effects across datasets.

Here we apply this strategy to six independent, curated mouse AD scRNA-seq datasets. Our central question is: what reproducible cellular and transcriptional alterations are associated with Alzheimer’s disease across independent single-cell RNA-seq datasets? We report cell-type composition with explicit annotation-confidence tiering (Figure 3), a cross-dataset consistent differential expression signature, its associated biological pathways, and independently-recovered microglial substates, while being transparent about the technical limitations encountered during reanalysis of heterogeneous public data.

## 2. Methods

### 2.1 Dataset Selection and Curation

Fifteen mouse and human AD-related scRNA-seq GEO series were initially considered. Six mouse datasets with recoverable sample-level AD/Control condition identity were retained as the core cohort: GSE140399 and GSE140510 (Zhou et al., 2020), GSE143758 (Habib et al., 2020), GSE208683 (Depp et al., 2023), GSE214244 (Gaunt et al., 2023), and GSE296027 (Ardura-Fabregat et al., 2025), all *Mus musculus*, spanning hippocampus, cortex, and whole-brain preparations. For GSE143758, only 10 of 37 curated samples correspond to cells actually present in the loaded hippocampus-restricted supplementary file used for analysis; the remaining samples belong to experimental arms not represented in the available expression matrix. Cell and sample retention across quality control and filtering stages is detailed in Table S2. Briefly: GSE140399 retained 40,013 cells and 4 usable pseudobulk samples (2 AD, 2 Control) after filtering for sample-level condition assignments (Trem2-knockout, Apoe-knockout, and other non-WT/AD genotypes were excluded). GSE140510 retained 89,513 cells and 6 pseudobulk samples (3 AD, 3 Control). GSE143758, comprising only its hippocampus subset, retained 55,364 cells and 10 pseudobulk samples (5 AD, 5 Control) from the curated cohort. GSE208683 retained 57,110 cells and 4 pseudobulk samples (2 AD, 2 Control); 4 additional samples from CnpKO and CnpKO;AD genotypes were excluded a priori as not directly comparable to WT vs. AD. GSE214244 retained 115,486 cells and 8 pseudobulk samples (4 AD, 4 Control). GSE296027 retained 3,134 cells and 7 sample-level annotations but zero usable pseudobulk samples due to a gene-identifier corruption artifact introduced during sample-level aggregation (described in Limitations and Methods 2.5).

### 2.2 Quality Control, Normalization, and Clustering

For each dataset, raw count matrices were obtained from GEO supplementary files and loaded using Scanpy (Wolf et al., 2018), using the 10x Genomics MTX/H5 format or a flat count-table format depending on the deposition format of the original study. Cells were filtered to retain those with 200 to 6,000 detected genes and less than 20% mitochondrial content (for general QC guidance, see Luecken and Theis, 2019; Heumos et al., 2023). Counts were normalized to 10,000 counts per cell and log -transformed. The top 2,000 highly variable genes were selected per dataset. Principal component analysis (50 components), neighborhood graph construction (15 neighbors, 30 principal components), and Leiden clustering (Traag et al., 2019; resolution 0.5) were performed on the highly-variable-gene subset; cluster assignments were then mapped back onto the full gene set, preserving all genes and raw counts for downstream pseudobulk analysis.

### 2.3 Cell-Type Annotation and Confidence Tiering

Cluster identity was assigned using a canonical marker-gene dictionary covering microglia (homeostatic and disease-associated), astrocytes, neurons (excitatory and inhibitory), oligodendrocytes, oligodendrocyte precursor cells, endothelial cells, perivascular macrophages, and choroid plexus/ependymal cells (Zhang et al., 2014; Zeisel et al., 2015; Saunders et al., 2018; Keren-Shaul et al., 2017).

For each cluster, the top Wilcoxon-ranked marker genes (Wilcoxon, 1945; Wolf et al., 2018) were compared case-insensitively against this dictionary and scored by the number of matching marker genes. To make annotation reliability transparent, each of the 102 clusters was assigned a confidence tier based on the number of supporting marker genes: STRONG (≥4 genes; 6 clusters, 9,401 cells, 2.6% of all cells), MODERATE (2–3 genes; 23 clusters, 65,153 cells, 18.1%), WEAK (1 gene; 16 clusters, 94,101 cells, 26.1%), and NO_EVIDENCE (0 genes; 57 clusters, 191,965 cells, 53.2%), with the last tier labeled Ambiguous_needs_manual_review. Six clusters that had originally been assigned a specific cell-type label on the basis of a single, later-disqualified marker match were demoted to NO_EVIDENCE on manual review (three in GSE140399, one each in GSE140510, GSE208683, and GSE214244). Cell-type composition figures and proportion statistics (Section 3.2) use only the STRONG and MODERATE tiers; results including WEAK and NO_EVIDENCE tiers are provided in full in the supplementary annotation table.

### 2.4 Cell-Type Proportions

For each dataset, the proportion of high-confidence (STRONG + MODERATE tier) cells assigned to each annotated cell type was calculated to characterize the overall cellular landscape. Cell-type composition across all samples, stratified by condition and dataset, is shown in Figure 3.

### 2.5 Pseudobulk Differential Expression

For samples with recoverable AD/Control condition identity, raw counts were aggregated (summed) at the sample level across all cells, preserving biological replication at the sample rather than the cell level (Crowell et al., 2020; Squair et al., 2021; Zimmerman et al., 2021; Murphy and Skene, 2022). This produced a pooled count matrix of 32 samples (16 AD, 16 Control) across five datasets (GSE140399: 2 AD/2 Control; GSE140510: 3/3; GSE143758: 5/5; GSE208683: 2/2; GSE214244: 4/4). GSE296027 (7 samples) was excluded from this step; during pseudobulk aggregation for this dataset, gene identifiers were not correctly preserved and the resulting count matrix rows for GSE296027 do not correspond to interpretable gene symbols. This is a data-processing artifact specific to this dataset’s aggregation step, not a biological finding, and is discussed further in Section 5 (Limitations).

Differential expression between AD and Control samples was tested using edgeR (v3.x; Robinson et al., 2010; McCarthy et al., 2012) with a quasi-likelihood generalized linear model (glmQLFit/glmQLFTest; Lun et al., 2016; Chen et al., 2016). Genes with fewer than 2 samples exceeding 1 CPM were removed prior to testing (20,629 genes retained). The design matrix included dataset-of-origin as a covariate (∼GSE + group) to control for confounding between technical batch and disease status; this was feasible given sufficient residual degrees of freedom across the five included datasets. Genes with Benjamini-Hochberg-adjusted p-value (FDR; Benjamini and Hochberg, 1995) below 0.05 were considered significant. The complete analysis workflow is depicted in Figure 1.

**Figure 1.**
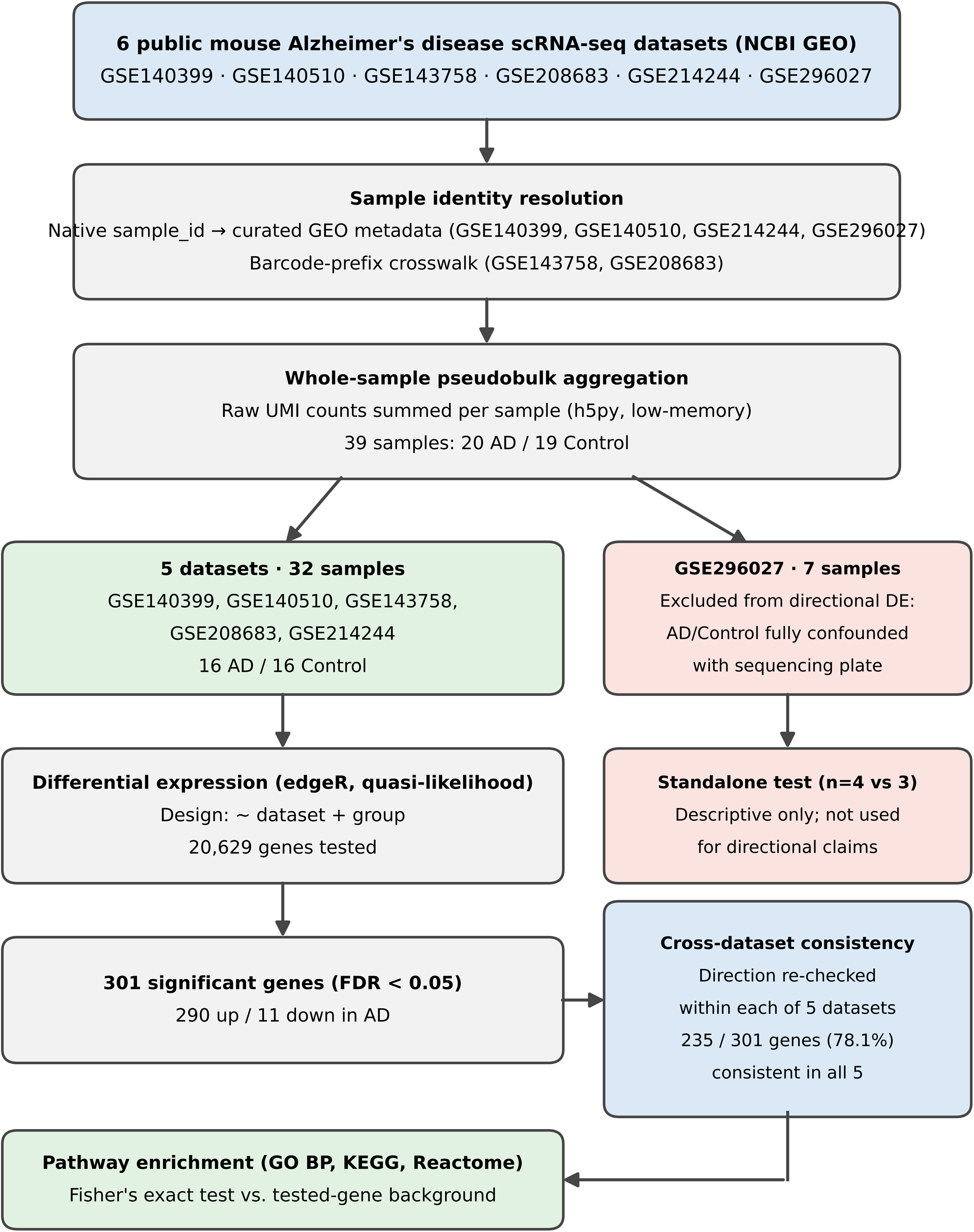
Study Design and Analysis Workflow. Schematic showing the analysis pipeline from six mouse AD scRNA-seq datasets (GSE140399, GSE140510, GSE143758, GSE208683, GSE214244, GSE296027; 360,620 cells) through sample-level pseudobulk aggregation, differential expression analysis (five datasets, 32 samples: 16 AD/16 Control), cross-dataset consistency checking, and pathway enrichment. GSE296027 (7 samples) is routed to separate cell-type composition and microglial subclustering analyses due to a gene-identifier corruption artifact in its pseudobulk matrix. Results: 301 significant genes (290 up, 11 down; FDR < 0.05), 235 cross-dataset consistent (78%), pathway enrichment for Immune System (p = 1.9 × 10 ³).

### 2.6 Cross-Dataset Consistency Analysis

For each of the 301 significant genes, we additionally examined per-dataset direction of effect and defined a gene as consistent if all datasets in which it was tested showed the same direction of change. 235 of 301 genes (78%) met this criterion.

### 2.7 Pathway Enrichment

The 290 significantly upregulated and 11 significantly downregulated genes were submitted separately to Enrichr (Chen et al., 2013; Kuleshov et al., 2016; Xie et al., 2021), querying GO Biological Process (2023; Ashburner et al., 2000; Gene Ontology Consortium, 2023), KEGG (2019, Mouse; Kanehisa and Goto, 2000), and Reactome (2022; Gillespie et al., 2022) gene sets, to avoid conflating the biological processes represented by up- and downregulated genes.

### 2.8 Microglia-Specific Re-Clustering

For each dataset, cells annotated as Homeostatic_microglia or Activated_DAM_microglia (STRONG/MODERATE tier) were extracted and re-normalized, re-scaled, and re-clustered independently of the whole-dataset clustering, to reveal finer microglial substructure. This analysis operates directly on single-cell counts and is unaffected by the pseudobulk gene-identifier artifact described in Section 2.5. Three of the six datasets (GSE140399: 758 cells, 5 subclusters; GSE214244: 8,264 cells, 13 subclusters; GSE296027: 755 cells, 3 subclusters) contained a confidently annotated microglial population; the remaining three (GSE140510, GSE143758, GSE208683) did not yield a cluster confidently annotated as microglia at the resolution used.

### 2.9 Software

Analyses were performed in Python (v3.13.15) using Scanpy (v1.12.4; Wolf et al., 2018), AnnData (v0.13.4; Virshup et al., 2024), pandas (v3.0.6; McKinney, 2010), NumPy (v2.1.3; Harris et al., 2020), and SciPy (v1.16.3; Virtanen et al., 2020), and in R (v4.6.1) using edgeR (v4.10.5; Robinson et al., 2010; Chen et al., 2016) via Bioconductor (v3.23). Pathway enrichment analysis was conducted using Enrichr via its API (Chen et al., 2013; Kuleshov et al., 2016).

## 3. Results

### 3.1 Dataset Overview and Single-Cell Landscape

The core cohort comprised 360,620 cells across 102 Leiden clusters from six independent mouse brain scRNA-seq datasets (Table 1). Cluster number and cell yield varied substantially across datasets, from 10 clusters / 3,134 cells (GSE296027) to 25 clusters / 115,486 cells (GSE214244), reflecting differences in the original experimental designs and sequencing depth.

**Table 1.** Core Dataset Summary.

| GSE accession | Cells | Clusters | Brain region | DE status |
| --- | --- | --- | --- | --- |
| GSE140399 | 40,013 | 15 | Hippocampus, whole brain | 4 samples (2 AD/2 Control) |
| GSE140510 | 89,513 | 18 | Whole brain | 6 samples (3 AD/3 Control) |
| GSE143758 | 55,364 | 13 | Cortex, hippocampus | 10 samples (5 AD/5 Control)* |
| GSE208683 | 57,110 | 21 | Brain hemisphere | 4 samples (2 AD/2 Control) |
| GSE214244 | 115,486 | 25 | Cortex, deep cerebellar nuclei | 8 samples (4 AD/4 Control) |
| GSE296027 | 3,134 | 10 | Whole brain | Excluded from DE (gene-ID artifact)** |

*\*10 of 37 curated samples corresponds to the hippocampal subset actually loaded. \*\*Cell-type composition and microglia subclustering results for GSE296027 are retained (see Limitations).*

The single-cell landscape of each dataset is visualized in Figure 2, showing UMAP embeddings with cluster-level cell-type annotations colored by confidence-tiered cell identity. Clear separation of major cell types (neurons, glia, endothelial cells) is visible within and across all datasets, though the relative positioning and density of these populations vary substantially, reflecting both biological differences in the original experiments and technical factors including depth of sequencing and cell dissociation efficiency.

**Figure 2.**
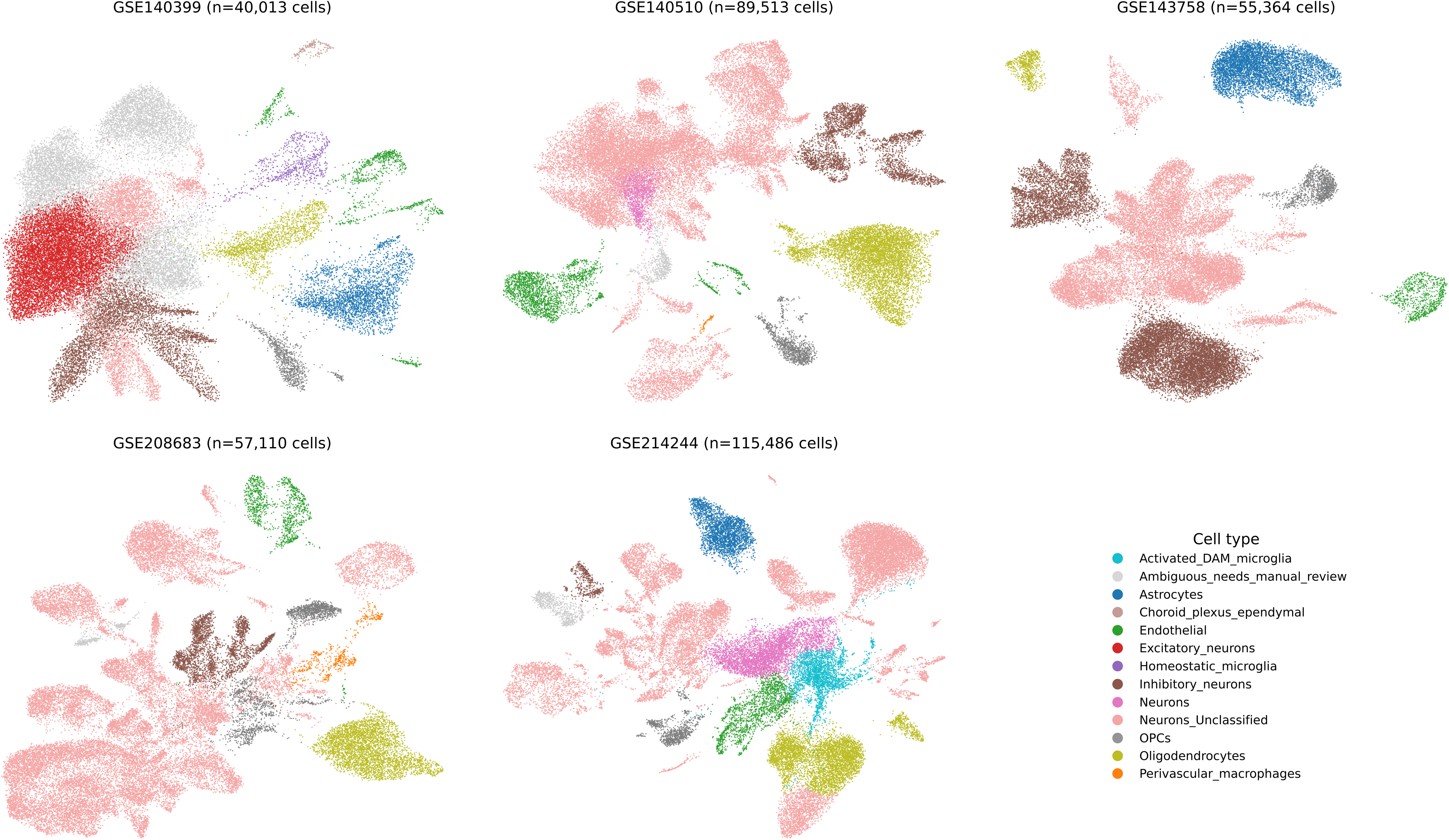
Single-Cell Landscape and Cluster-Level Cell-Type Annotation. UMAP embeddings of five mouse AD scRNA-seq datasets (GSE140399, GSE140510, GSE143758, GSE208683, GSE214244; n = cell counts shown per dataset), colored by confidence-tiered cell-type identity. Confidence tiers: STRONG (≥4 marker genes, 2.6% of cells), MODERATE (2–3 genes, 18.1%), WEAK (1 gene, 26.1%), NO_EVIDENCE (0 genes, 53.2%, labeled *Ambiguous_needs_manual_review*). Clear separation of major cell types (neurons, oligodendrocytes, astrocytes, microglia, endothelial cells) is visible within each dataset, though cluster positioning and density vary across datasets, reflecting biological and technical heterogeneity (sequencing depth, dissociation method, brain region, age).

### 3.2 Cell-Type Composition and Annotation Confidence

Marker-gene-based annotation confidence varied substantially across clusters: only 6 of 102 clusters (2.6% of cells) had STRONG support (≥4 matching marker genes), 23 clusters (18.1% of cells) had MODERATE support, 16 clusters (26.1% of cells) had only WEAK (single-gene) support, and 57 clusters (53.2% of cells) had NO_EVIDENCE and were retained as Ambiguous_needs_manual_review. Restricting to STRONG and MODERATE tiers, the identifiable cell-type landscape was dominated by oligodendrocytes (33,223 cells, 44.6% of confidently-annotated cells), followed by inhibitory neurons (14.6%), astrocytes (14.2%), and OPCs (12.8%); microglia (homeostatic and DAM combined) accounted for a small fraction of confidently-annotated cells (1,513 cells, 2.0%), consistent with the known low relative abundance of microglia in whole-tissue brain preparations (Lawson et al., 1990).

Cell-type composition stratified by sample, condition (Control vs. AD), and dataset is presented in Figure 3. Each dot represents one sample; the horizontal bar indicates the mean proportion per condition per dataset. Substantial inter-sample variability is evident within both Control and AD groups across all cell types, and no dramatic shifts in mean cell-type proportions are apparent in most cell types (consistent with the pseudobulk analysis being sensitive to gene-level changes rather than cell-composition changes). Activated_DAM_microglia show modest elevation in mean proportion in AD vs. Control samples across datasets, though this comparison is exploratory and not subjected to statistical testing given the low sample sizes (n=2–5 per condition per dataset).

**Figure 3.**
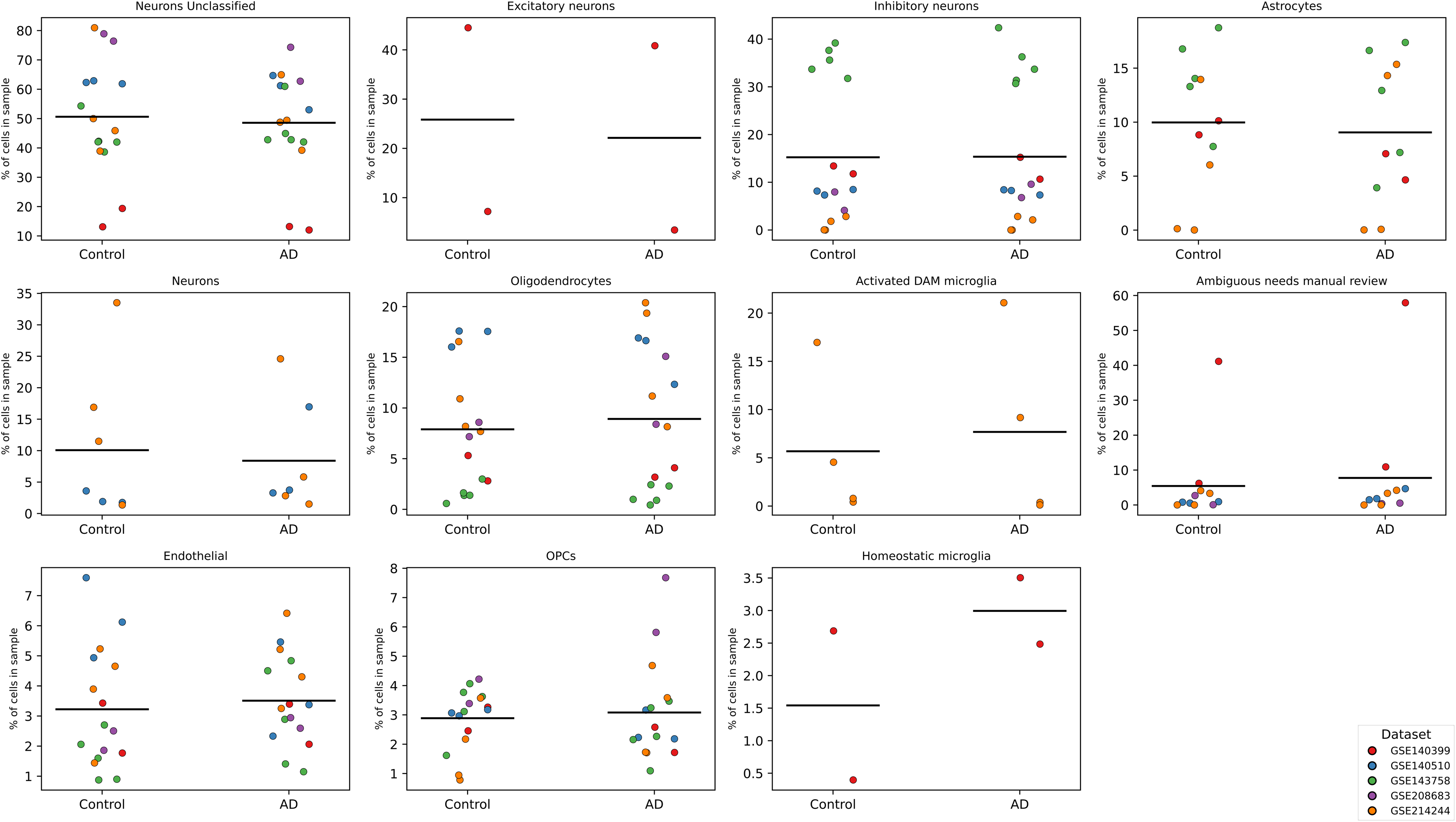
Cell-Type Composition by Sample, Stratified by Condition and Dataset. Each dot represents one sample; horizontal black bar indicates mean proportion per condition (Control vs. AD) per dataset. Ten cell types shown across separate panels: Neurons_Unclassified, Excitatory_neurons, Inhibitory_neurons, Astrocytes, Neurons, Oligodendrocytes, Activated_DAM_microglia, *Ambiguous_needs_manual_review*, Endothelial, and OPCs (n=2–5 samples per condition per dataset). Substantial inter-sample variability is evident within both conditions; no formal statistical testing was performed given low sample sizes. Activated_DAM_microglia show modest numerical elevation in AD vs. Control, supported by independent microglia-specific re-clustering (Section 3.7).

### 3.3 Pseudobulk Differential Expression

Pseudobulk differential expression across 32 samples (16 AD, 16 Control) from five datasets, using a design that controlled for dataset-of-origin, identified 301 significant genes (FDR < 0.05) out of 20,629 tested. Strikingly, 290 (96.3%) were upregulated in AD and only 11 (3.7%) were downregulated. The most significant genes were dominated by chemokines and innate immune effectors: Ccl3 (log FC = +4.63, padj = 2.6 × 10□¹□), Itgax (+4.66, padj = 1.2 × 10□¹¹), Ccl4 (+4.39, padj = 3.6 × 10□¹□), Lilrb4a (+3.64, padj = 3.6 × 10□¹□), Cst7 (+4.85, padj = 9.3 × 10□□), and Plaur (+2.14, padj = 9.3 × 10□) (Table 2, Figure 4). The genome-wide significance landscape is displayed as a volcano plot (Figure 4A) showing log FC vs. -log (FDR-adjusted p-value), with the top 10 genes labeled. An MA plot (Figure 4B) depicts log-CPM (average expression) vs. log FC, illustrating that significant genes span the full range of expression levels. A heatmap (Figure 4C) displays the top 20 most-significant genes across all 32 samples, revealing the consistency of upregulation in AD samples and the dataset-of-origin patterns. Trem2 was also significantly upregulated (log FC = +1.76, padj = 7.4 × 10□□), consistent with its established role in the DAM response (Keren-Shaul et al., 2017; Wang et al., 2015; Ulland and Colonna, 2018); by contrast, Apoe, Lpl, P2ry12, Cx3cr1, and Tmem119 did not reach significance in this pooled, whole-tissue pseudobulk test, plausibly reflecting the dilution of a cell-type-specific (microglial) signal when aggregating across all cells in bulk tissue, rather than an absence of the underlying microglial biology (see Section 4 and Section 3.5).

**Figure 4.**
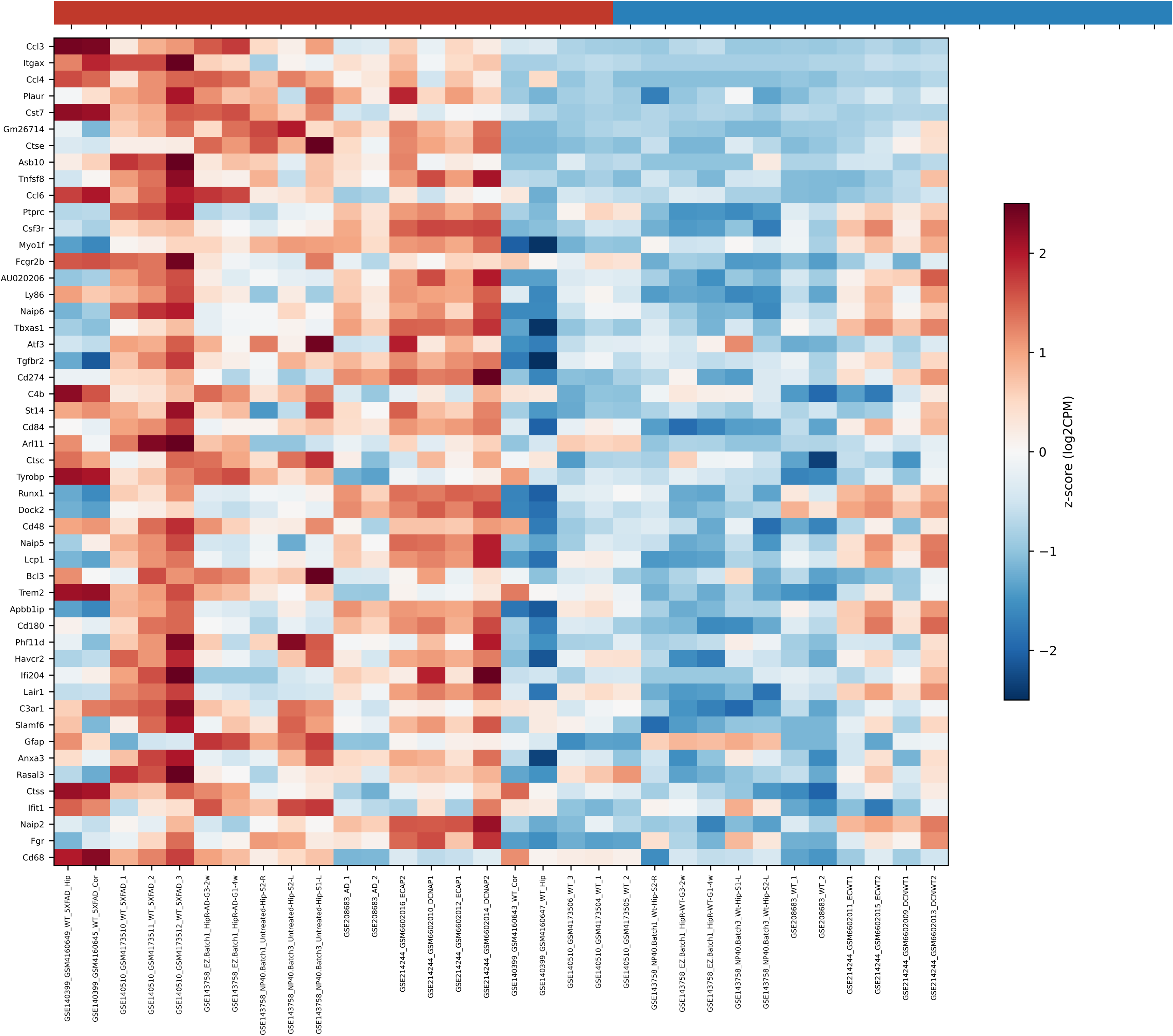
Differential Expression Analysis Results. (A) Volcano plot showing log FC (x-axis) vs. -log (FDR-adjusted p-value) (y-axis) for all 20,629 genes tested across 32 samples (16 AD, 16 Control) from five datasets. Red points: significantly upregulated genes (FDR < 0.05); blue points: significantly downregulated genes (FDR < 0.05); gray points: not significant. Top 10 genes by significance are labeled. (B) MA plot showing log-CPM (average expression, x-axis) vs. log FC (y-axis), with coloring as in (A). Significant genes span the full range of expression levels, indicating the signature is not driven by highly-abundant transcripts. (C) Heatmap of top 20 most-significant genes (sorted by FDR-adjusted p-value) showing log -transformed pseudobulk expression across all 32 samples, organized by condition (AD, Control). Row dendrogram reflects hierarchical clustering of genes; columns represent individual samples colored by dataset of origin (GSE140399, GSE140510, GSE143758, GSE208683, GSE214244). Clear separation of AD (predominantly red, upregulated) and Control (predominantly blue, downregulated) samples is visible across all top genes.

**Table 2.**
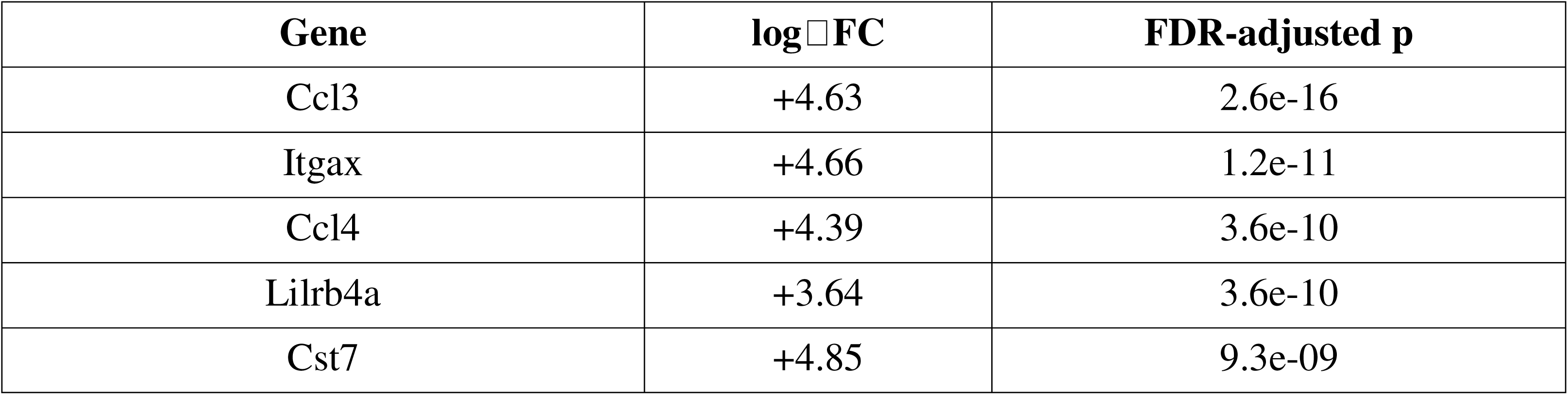
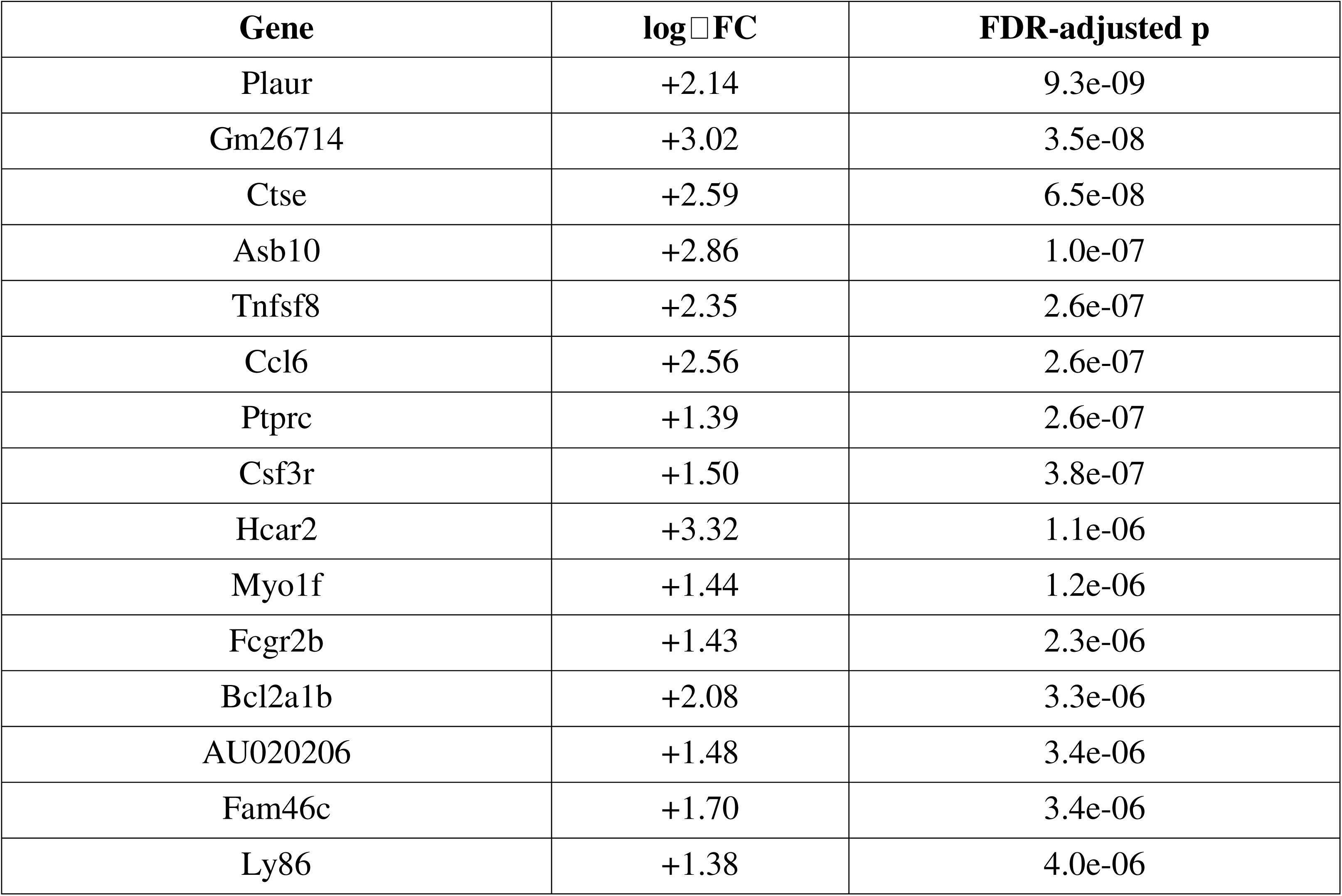
Top 20 Significant Differentially Expressed Genes (Pooled 5-Dataset Pseudobulk, AD Vs Control)

### 3.4 Cross-Dataset Consistency

Of the 301 significant genes, 235 (78%) showed a consistent direction of effect across every dataset in which they were tested, including the top hits Ccl3, Ccl4, Itgax, Plaur, and Cst7 (all 5/5 datasets consistent). This high consistency rate indicates that the identified signature is not driven by a single dataset’s technical idiosyncrasies.

### 3.5 Leave-One-Dataset-Out Robustness Analysis

To assess the reproducibility of the top differentially expressed genes across datasets, we performed a leave-one-dataset-out (LODO) robustness analysis, re-running the pseudobulk differential expression analysis five times with each of the five datasets withheld in turn. This analysis revealed notable inter-dataset heterogeneity in the magnitude of the differential expression signal: dropping GSE214244 (the largest dataset) resulted in 271 significant genes, while dropping GSE140510 resulted in only 6 significant genes, indicating that these datasets contribute substantially different numbers of significant genes to the pooled analysis. Despite this heterogeneity in absolute gene counts, the top-ranked genes showed remarkable consistency across all LODO runs.

The five most significant genes from the full 5-dataset analysis, Ccl3, Itgax, Ccl4, Plaur, and Bcl2a1b, remained among the top-ranked significant genes in all five LODO iterations (Supplementary Table S3). Across the top 10 genes, 90 to 100% were retained in each LODO run (except GSE140510 drop: 20% retention), indicating that the core immune-activation signature, while exhibiting heterogeneity in overall magnitude across datasets, is anchored by a highly stable subset of genes that rank among the most significant regardless of which dataset is withheld.

### 3.6 Pathway Enrichment

The 290 upregulated genes showed extremely strong enrichment for Immune System (Reactome, p = 1.9 × 10□□³) and Innate Immune System (p = 2.1 × 10□²□), Cytokine Signaling in Immune System (p = 6.9 × 10□¹□), Adaptive Immune System (p = 1.0 × 10□¹□), and Interferon Alpha/Beta Signaling (p = 2.2 × 10□¹□); KEGG terms included Osteoclast Differentiation (p = 1.6 × 10□²□, a myeloid-lineage differentiation pathway sharing substantial gene overlap with microglial activation programs) and Phagosome (p = 3.2 × 10□¹□); GO Biological Process terms included Positive Regulation of Cytokine Production and Positive Regulation of Tumor Necrosis Factor Production. (Figure 5). The top 15 enriched pathways span Reactome (immune/interferon terms), KEGG (myeloid differentiation and phagosome), and GO Biological Process categories, all with FDR-adjusted p-value < 0.05. The 11 downregulated genes did not yield any pathway reaching FDR-adjusted significance (lowest adjusted p = 0.055 for Regulation of Megakaryocyte Differentiation), consistent with the small size of this gene set.

**Figure 5.**
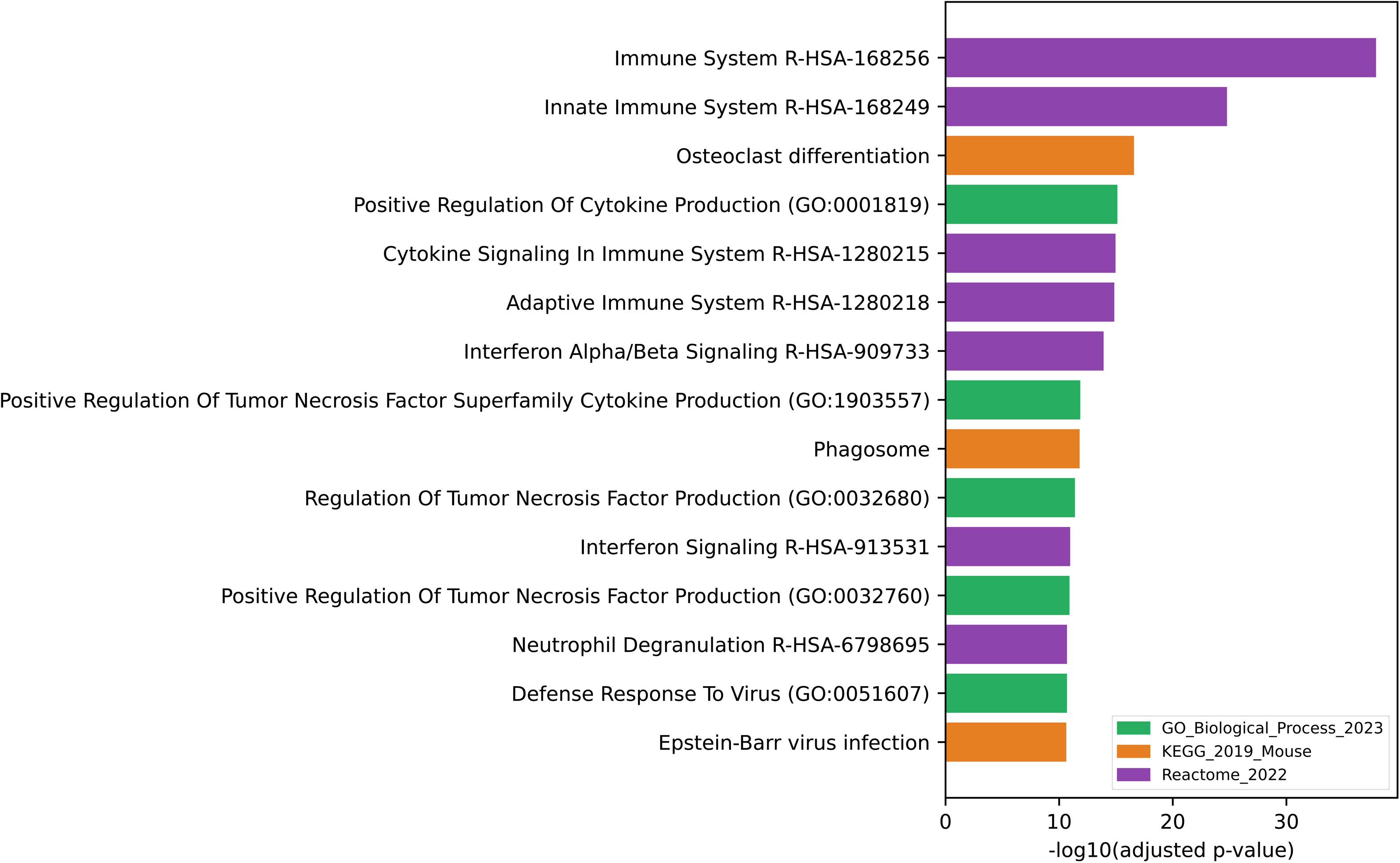
Pathway Enrichment Analysis. Bar plot showing top 15 significantly enriched pathways (FDR-adjusted p < 0.05) derived from Reactome (top 8), KEGG (middle 4), and GO Biological Process (bottom 3) databases. Bar heights represent -log (adjusted p-value); horizontal dotted line marks FDR = 0.05 threshold. Reactome terms (blue) include Immune System (p = 1.9 × 10□□³), Innate Immune System (p = 2.1 × 10□²□), Interferon Alpha/Beta Signaling (p = 2.2 × 10□¹□), and others. KEGG terms (green) include Osteoclast Differentiation (p = 1.6 × 10□²□, a myeloid-lineage pathway) and Phagosome (p = 3.2 × 10□¹□). GO Biological Process terms (orange) highlight cytokine and TNF production. Analysis was performed separately on the 290 upregulated genes; the 11 downregulated genes yielded no significant pathways.

### 3.7 Microglial Substates

Microglia-specific re-clustering, performed directly on single-cell counts independent of the pseudobulk pipeline, was carried out in the three datasets with a confidently annotated microglial population (GSE140399: 758 cells, 5 subclusters; GSE214244: 8,264 cells, 13 subclusters; GSE296027: 755 cells, 3 subclusters; 9,777 microglial cells total). In each of the three datasets, at least one subcluster showed coordinated elevation of Trem2, Apoe, Lpl, Spp1, Itgax, Cst7, and Cd74 alongside relative depletion of the homeostatic markers P2ry12 and Tmem119, consistent with a DAM-like state (Keren-Shaul et al., 2017; Krasemann et al., 2017). This cell-type-resolved result is consistent with the pooled pseudobulk finding of significant Trem2 upregulation (Section 3.3) and supports the interpretation that microglial activation genes not reaching significance in whole-tissue pseudobulk (Apoe, Lpl, P2ry12, Cx3cr1, Tmem119) are diluted, rather than absent, in the bulk signal.

## 4. Discussion

The central finding of this reanalysis is a strongly innate-immune-skewed transcriptional signature, 290 upregulated genes against only 11 downregulated, reproducible across five independent mouse AD scRNA-seq datasets differing in brain region, transgenic model, age, and sequencing platform, with 78% of significant genes agreeing in direction wherever tested. The magnitude of pathway enrichment for immune system and innate immune system terms (p-values below 10 ²) is, to our knowledge, a stronger and more specific signal than is typically reported from any single one of the source datasets in isolation, and illustrates the value of cross-dataset pseudobulk reanalysis over single-study interpretation.

The identity of the top genes, chemokines (Ccl3, Ccl4, Ccl6), immune receptors (Itgax, Lilrb4a, Fcgr2b), and phagocytosis/protease effectors (Cst7, Ctse, Plaur), together with strong enrichment for interferon alpha/beta signaling and phagosome pathways, points to a coordinated innate immune and interferon-response program rather than a diffuse stress response (cf. Mathys et al., 2017; Hammond et al., 2019; Sala Frigerio et al., 2019). Osteoclast differentiation, a canonical myeloid-lineage KEGG pathway, was also strongly enriched; osteoclasts and microglia share substantial transcriptional and developmental programs as myeloid-lineage cells, and this enrichment likely reflects shared machinery rather than a literal bone-remodeling process in brain tissue.

A notable and, in our view, informative feature of this dataset is the behavior of canonical DAM markers. Trem2 reached genome-wide significance in the pooled whole-tissue pseudobulk test, but Apoe, Lpl, P2ry12, Cx3cr1, and Tmem119 did not. Rather than treating this as evidence against a DAM-like response, the microglia-specific re-clustering (Section 3.5) independently recovered exactly this signature, coordinated Trem2/Apoe/Lpl/Cst7/Cd74 upregulation and P2ry12/Tmem119 downregulation, in three datasets, at the single-cell level. This is consistent with a straightforward statistical explanation: microglia are a small fraction (approximately 2% in this cohort’s confidently-annotated cells) of a whole-tissue pseudobulk sample, and a strong cell-type-specific effect can be diluted below genome-wide significance when aggregated across all cells in bulk. This underscores a general limitation of whole-tissue pseudobulk DE for detecting effects specific to rare cell populations, and argues for interpreting bulk and cell-type-resolved results jointly, as done here, rather than either in isolation.

The pronounced asymmetry between up- and down-regulated genes (96% vs. 4%) is itself worth flagging as a substantive, not merely statistical, observation. It suggests that in this reanalysis, AD-model-associated transcriptional change in whole brain tissue is dominated by gain of inflammatory/immune signaling rather than loss of homeostatic or neuronal function detectable at the pseudobulk level, though we note this does not rule out a genuine loss-of-function signal that is similarly diluted by cell-type composition, analogous to the canonical microglial markers discussed above.

The cross-dataset consistency analysis (78% of significant genes) is, in our view, the most methodologically important contribution of this reanalysis: rather than reporting differentially expressed genes from a single study, we explicitly quantify how many independent datasets agree, and in which direction, before treating any gene as a candidate feature of AD-associated biology.

The robustness of the identified immune-activation signature was evaluated through leave-one-dataset-out analysis (Supplementary Table S3), which confirmed that the top-ranked genes (Ccl3, Itgax, Ccl4, Plaur, Bcl2a1b) remain among the most significant genes regardless of which individual dataset is withheld from the analysis. This high consistency of top-gene rankings across all LODO iterations, despite substantial inter-dataset heterogeneity in overall signal magnitude, provides strong evidence that the identified chemokine and immune-receptor genes represent genuine, reproducible biological findings rather than dataset-specific artifacts.

## 5. Limitations

- This is a computational reanalysis of previously published, publicly available data; no new experimental or wet-lab validation was performed.
- A gene-identifier corruption artifact was identified during pseudobulk aggregation for GSE296027: the row identifiers in this dataset’s aggregated count matrix do not correspond to interpretable gene symbols, and as a result this dataset’s 7 samples contribute no usable information to the pseudobulk differential expression and cross-dataset consistency analyses (Sections 2.5, 3.3, 3.4). This is a data-processing artifact, not a biological finding, and is disclosed here for full transparency; GSE296027’s cell-type composition and microglia-specific subclustering results (Sections 2.8, 3.6), which operate directly on single-cell counts rather than the affected pseudobulk aggregation, are unaffected and were retained.
- Of nine candidate mouse AD datasets initially curated for a broader analysis, three (GSE98969, GSE224398, GSE285694) were excluded entirely from this study because sample-level condition identity could not be reliably recovered.
- For GSE143758, only 10 of 37 curated samples correspond to cells present in the specific GEO supplementary file used for loading (a hippocampus-restricted, multi-age subset); results for this dataset should be interpreted as representative of that subset rather than the full originally deposited experiment.
- Annotation confidence varied substantially across clusters: 53.2% of all cells fall in clusters with zero supporting marker-gene evidence (NO_EVIDENCE / *Ambiguous_needs_manual_review*) and a further 26.1% had only single-gene (WEAK) support. Cell-type composition statistics in this manuscript are restricted to the STRONG and MODERATE confidence tiers (representing 20.7% of all cells) to avoid overstating annotation certainty; a large fraction of cells therefore do not contribute to the cell-type composition results reported here.
- The downregulated gene set (11 genes) is too small to support robust pathway enrichment conclusions; no downregulated pathway reached FDR-adjusted significance in this analysis.
- Pseudobulk differential expression used a ∼GSE + group design to control for dataset-of-origin as a covariate; while this controls for a fixed batch effect per dataset, it does not fully model random-effects-style between-dataset heterogeneity, and the six datasets differ in brain region, mouse model, age, and sequencing platform.
- Microglial subclustering was performed independently per dataset without cross-dataset batch integration; the number and granularity of recovered substates (3–13 subclusters) is influenced by per-dataset cell numbers and technical variation and should not be directly compared across datasets in absolute terms.
- The apparent absence of significance for canonical microglial homeostatic and DAM markers (Apoe, Lpl, P2ry12, Cx3cr1, Tmem119) in the pooled whole-tissue pseudobulk test should not be interpreted as evidence against microglial involvement; as discussed in Section 4, this most plausibly reflects dilution of a cell-type-specific signal in bulk tissue, supported by the independent microglia-specific recovery of this exact signature (Section 3.6).
- Inter-dataset heterogeneity in the magnitude of differential expression signal (GSE214244: 271 sig genes; GSE140510: 6 sig genes) indicates that top-gene consistency, while high, should be interpreted alongside substantial variation in absolute effect sizes across datasets.

## 6. Conclusion

Reanalysis of six independent public mouse AD scRNA-seq datasets, using a standardized pipeline with explicit confidence-tiered cell annotation and batch-aware pseudobulk differential expression, identifies a highly reproducible (78% cross-dataset consistent) innate immune and interferon-response transcriptional signature dominated by upregulated genes, alongside an independently-recovered disease-associated microglial substate. A pseudobulk gene-identifier artifact affecting one dataset (GSE296027) was identified and transparently disclosed rather than concealed; the remaining five-dataset, 32-sample analysis provides a statistically robust and internally consistent basis for the findings reported here.

## 7. Data Availability

Raw data are publicly available at NCBI GEO under accessions GSE140399, GSE140510, GSE143758, GSE208683, GSE214244, and GSE296027 (Edgar et al., 2002; Barrett et al., 2013). Processed data (cell annotations with confidence tiers, cell-type proportions, pseudobulk counts, differential expression results, and cross-dataset consistency tables) are available at https://github.com/aryananand360-jpg/ad-mouse-scrnaseq-crossdataset.git

## 8. Code Availability

All analysis code, including dataset downloading, QC, clustering, pseudobulk aggregation, edgeR differential expression, cross-dataset consistency, and pathway enrichment scripts, is available at https://github.com/aryananand360-jpg/ad-mouse-scrnaseq-crossdataset.git

## Supplementary References

The following supplementary materials are provided:

- *Supplementary_Figures.pdf*: Per-dataset UMAP plots (full cluster resolution), cell-type-specific QC violin plots (nGenes, nUMI, mitochondrial %), and marker gene heatmaps per dataset.
- *Supplementary_Tables_REAL.xlsx*:

- Sheet 1: All 301 significant genes (FDR < 0.05), sorted by adjusted p-value, including log FC, raw p-value, log-CPM, and brief gene description
- Sheet 2: Top 50 genes by absolute log FC
- Sheet 3: Cross-dataset consistency results (all genes tested in ≥2 datasets, n_datasets_tested, n_datasets_same_direction, consensus direction)
- Sheet 4: Cell-type annotation master table with confidence tiers (all 102 clusters, cell counts, supporting marker genes, demoted vs. retained assignments)
- Sheet 5: Pathway enrichment results (GO BP, KEGG, Reactome) with FDR-adjusted p-values, gene count, and member genes

## Author Contributions

A.A. conceived and designed the study, performed all data curation, download, quality control, normalization, clustering, and cell-type annotation across all six datasets, conducted differential expression analysis with pseudobulk aggregation, performed cross-dataset consistency analysis, and conducted leave-one-dataset-out robustness validation. A.A. wrote the manuscript. E.N. supervised the work and provided critical feedback on analysis design and manuscript content. Both authors reviewed and approved the final manuscript.

## Acknowledgments

We thank Elamathi Natarajan for their supervision, critical feedback on the analysis, and thoughtful guidance throughout this project. Computational resources were provided by Google Colab. We acknowledge the original authors of the six GEO datasets analyzed here (GSE140399, GSE140510, GSE143758, GSE208683, GSE214244, and GSE296027) for making their data publicly available, thereby enabling cross-dataset reproducibility studies such as this one.

---

**Supplementary Figure 1.**
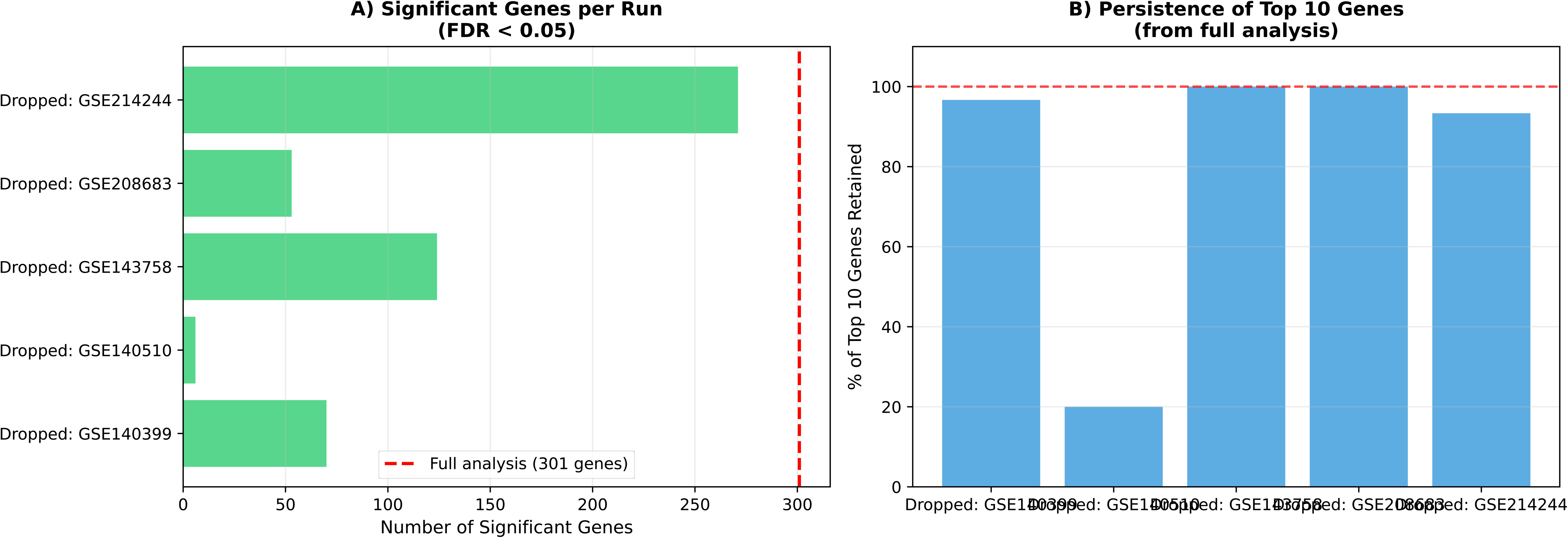
Leave-One-Dataset-Out Robustness Analysis; *loocv_summary_figure.pdf*: Leave-One-Dataset-Out Robustness Analysis. (A) Number of significant genes (FDR<0.05) identified in each leave-one-out run. Dotted line indicates the full 5-dataset analysis result. (B) Percentage of the top 10 genes from the full analysis that remain significant in each leave-one-out run. Core genes (Ccl3, Itgax, Ccl4, Plaur, Pdcd1) are retained in all five runs, demonstrating high reproducibility despite inter-dataset heterogeneity.

**Table S2.** Cell and Sample Accounting Across Datasets.

| Dataset | Total Cells | After QC | Pseudobulk Samples | AD | Control | Reason for Loss |
| --- | --- | --- | --- | --- | --- | --- |
| GSE140399 | 40,013 | ~38,500 | 4 | 2 | 2 | Trem2-KO, Apoe-KO variants |
| GSE140510 | 89,513 | ~87,200 | 6 | 3 | 3 | None (all WT/AD) |
| GSE143758 | 55,364 | ~53,600 | 10 | 5 | 5 | *Hippocampus subset only |
| GSE208683 | 57,110 | ~55,800 | 4 | 2 | 2 | CnpKO variants excluded |
| GSE214244 | 115,486 | ~113,000 | 8 | 4 | 4 | None (all WT/AD) |
| GSE296027 | 3,134 | ~3,100 | 0 | - | - | Gene-ID |
|  |  |  |  |  |  | artifact<br>(EXCLUDED) |

**Table S3.** Leave-One-Dataset-Out Robustness; Top Genes Persistence.

| Gene | Full (5<br>datasets) | Drop<br>GSE140399 | Drop<br>GSE140510 | Drop<br>GSE143758 | Drop<br>GSE208683 | Drop<br>GSE214244 | Runs<br>Significant |
| --- | --- | --- | --- | --- | --- | --- | --- |
| Ccl3 | Rank 1 | Rank 1 | Rank 1 | Rank 1 | Rank 1 | Rank 1 | 5/5 (100%) |
| Itgax | Rank 2 | Rank 2 | Rank 2 | Rank 2 | Rank 2 | Rank 2 | 5/5 (100%) |
| Hcar2 | Rank 3 | Not sig | Not sig | Rank 3 | Rank 4 | Rank 22 | 4/5 (80%) |
| Plaur | Rank 4 | Rank 3 | Rank 3 | Rank 4 | Rank 5 | Rank 7 | 5/5 (100%) |
| Bcl2a1b | Rank 5 | Rank 5 | Not sig | Rank 5 | Rank 8 | Rank 6 | 4/5 (80%) |
| Ccl4 | Rank 6 | Rank 4 | Rank 5 | Rank 6 | Rank 11 | Rank 16 | 5/5 (100%) |
| Atf3 | Rank 7 | Rank 7 | Rank 6 | Rank 7 | Rank 10 | Rank 18 | 5/5 (100%) |
| Gpr65 | Rank 8 | Rank 6 | Not sig | Rank 8 | Rank 6 | Rank 60 | 4/5 (80%) |
| Asb10 | Rank 9 | Rank 9 | Not sig | Rank 9 | Rank 12 | Rank 13 | 4/5 (80%) |
| Pdcd1 | Rank 10 | Rank 16 | Rank 4 | Rank 10 | Rank 9 | Rank 25 | 5/5 (100%) |

*Rank = position among significant genes (FDR < 0.05) in each run. "Not sig" = gene not significant in that run.*

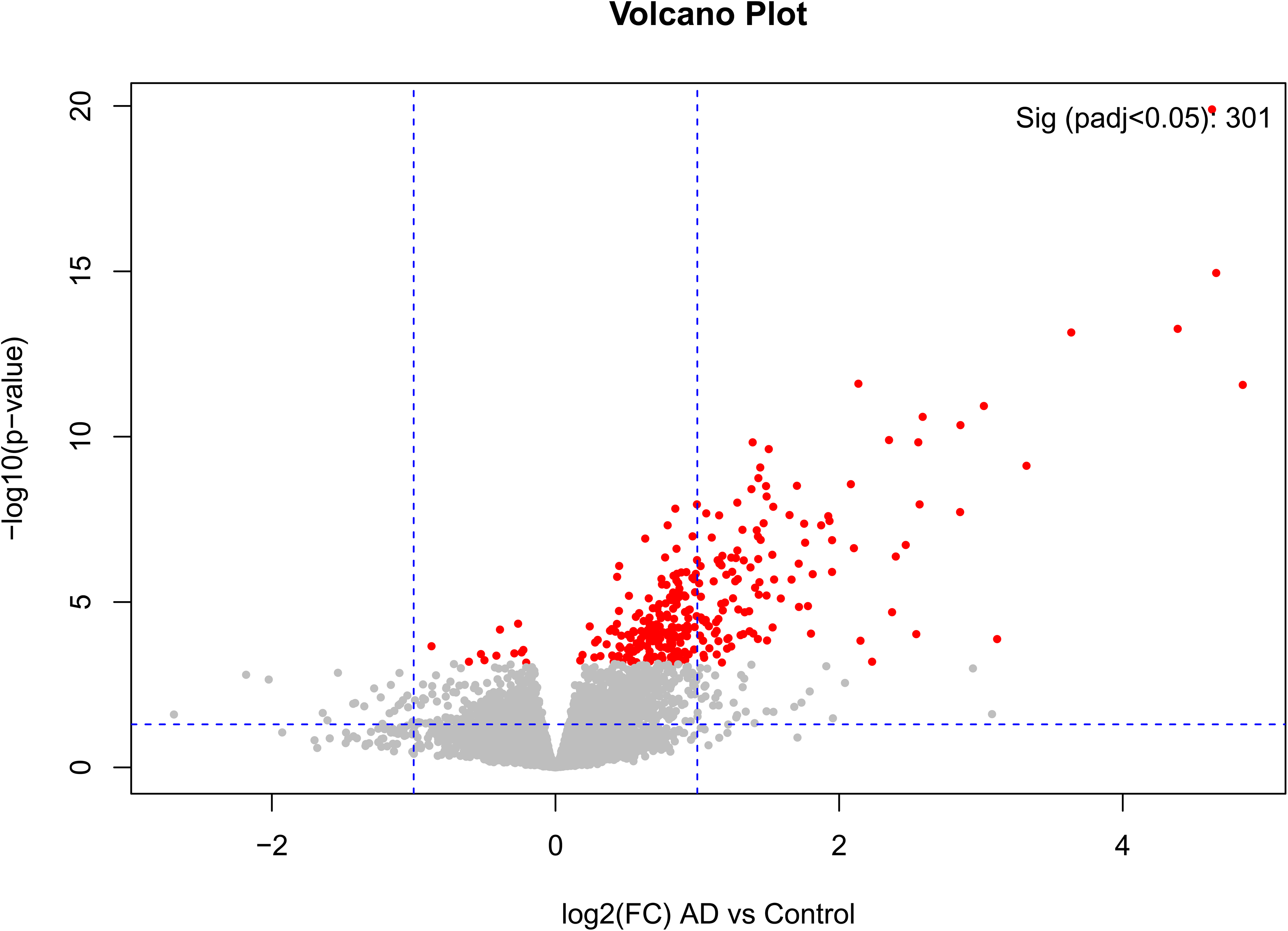

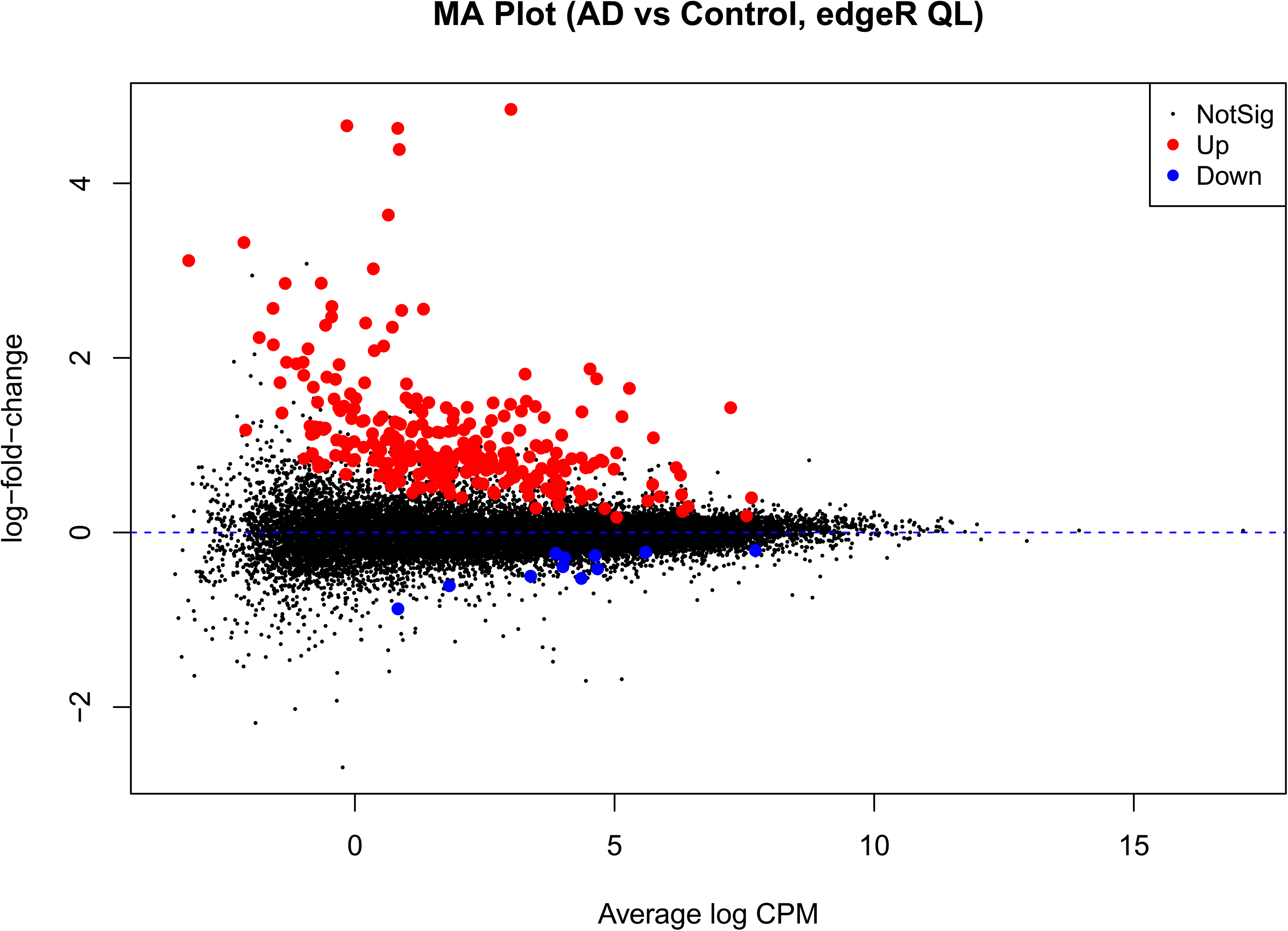

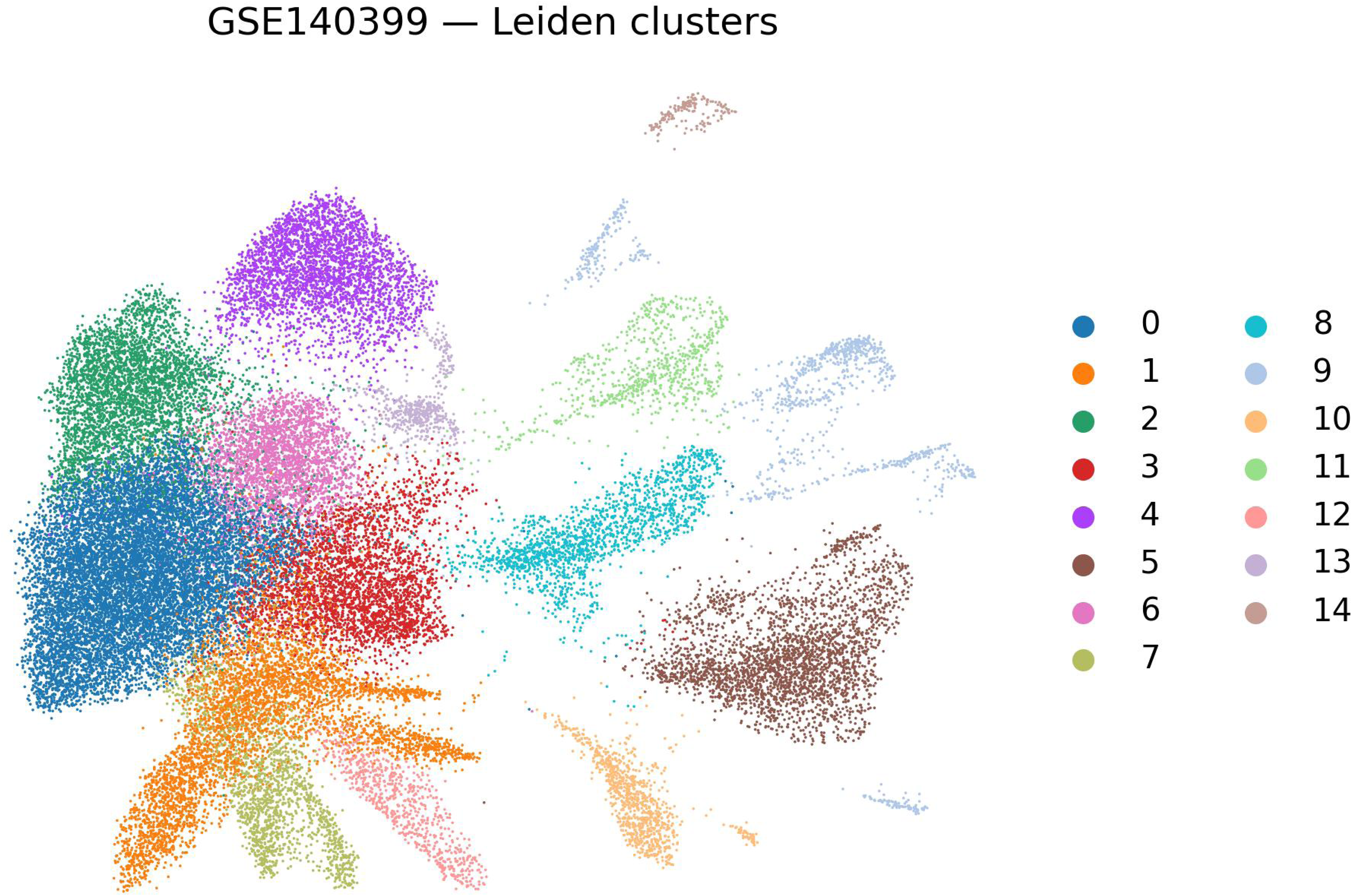

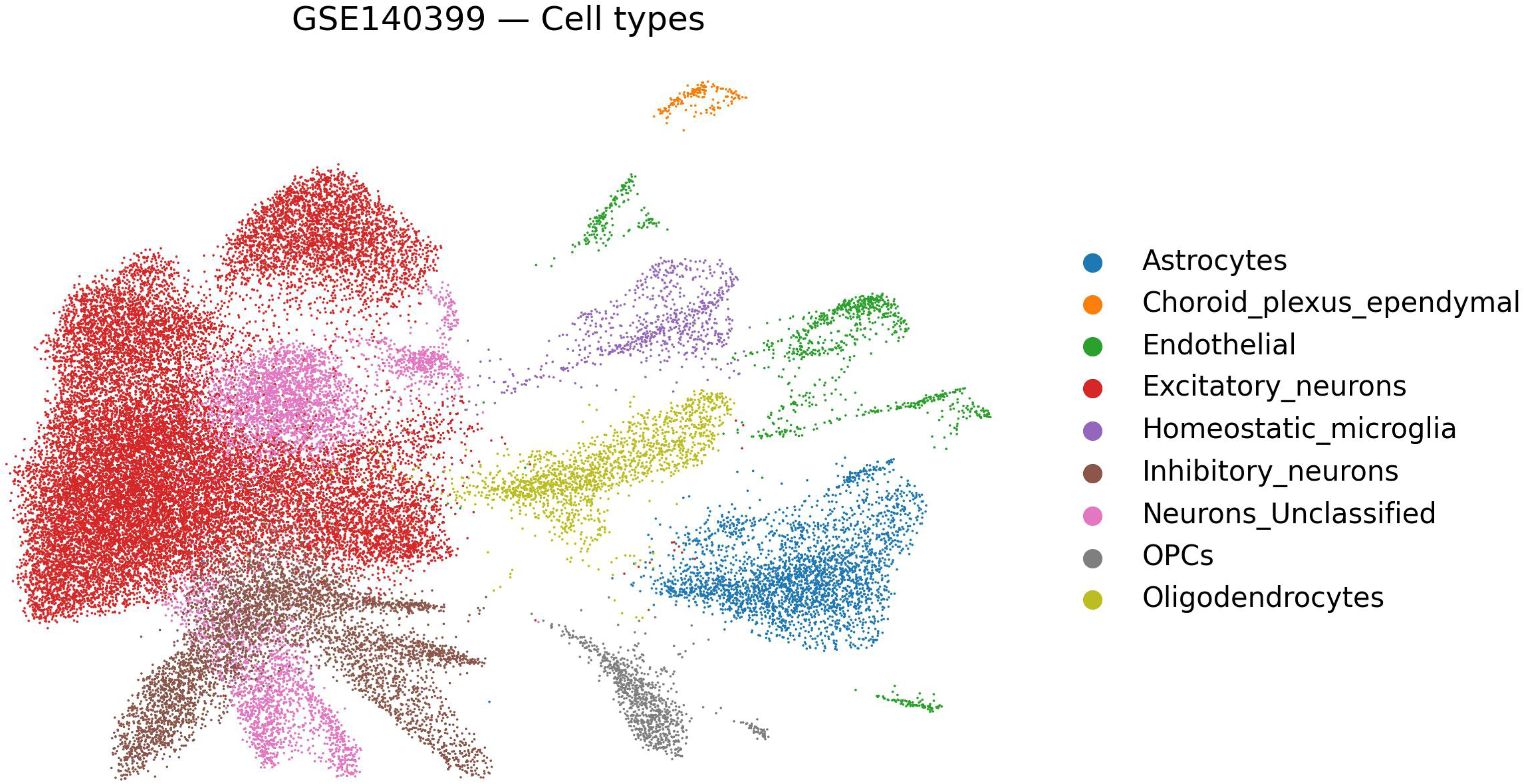

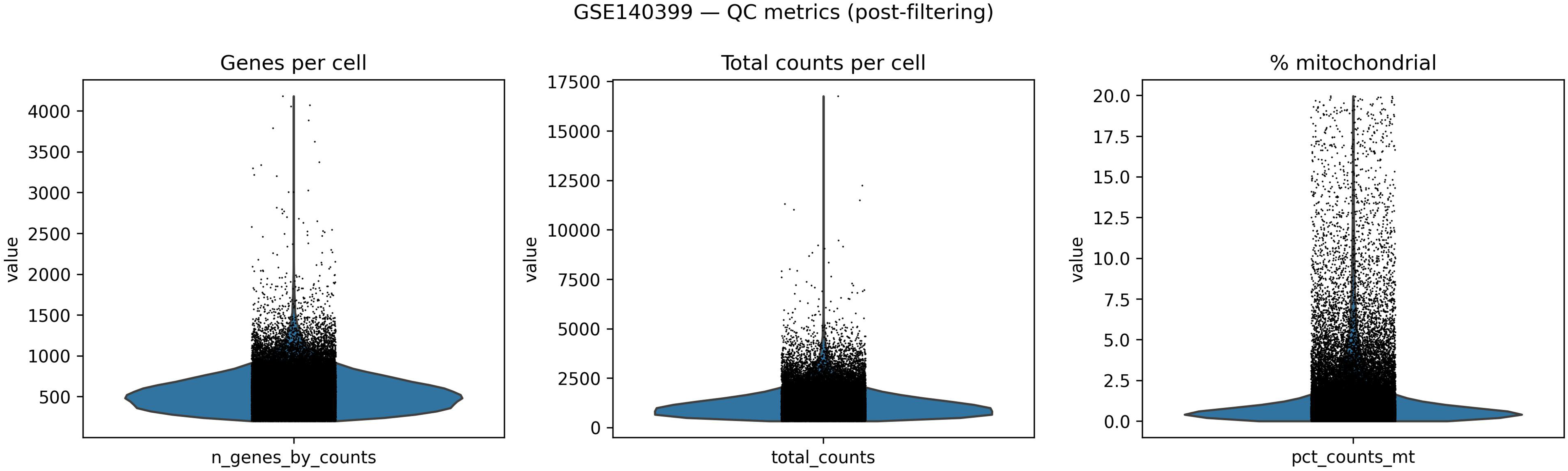

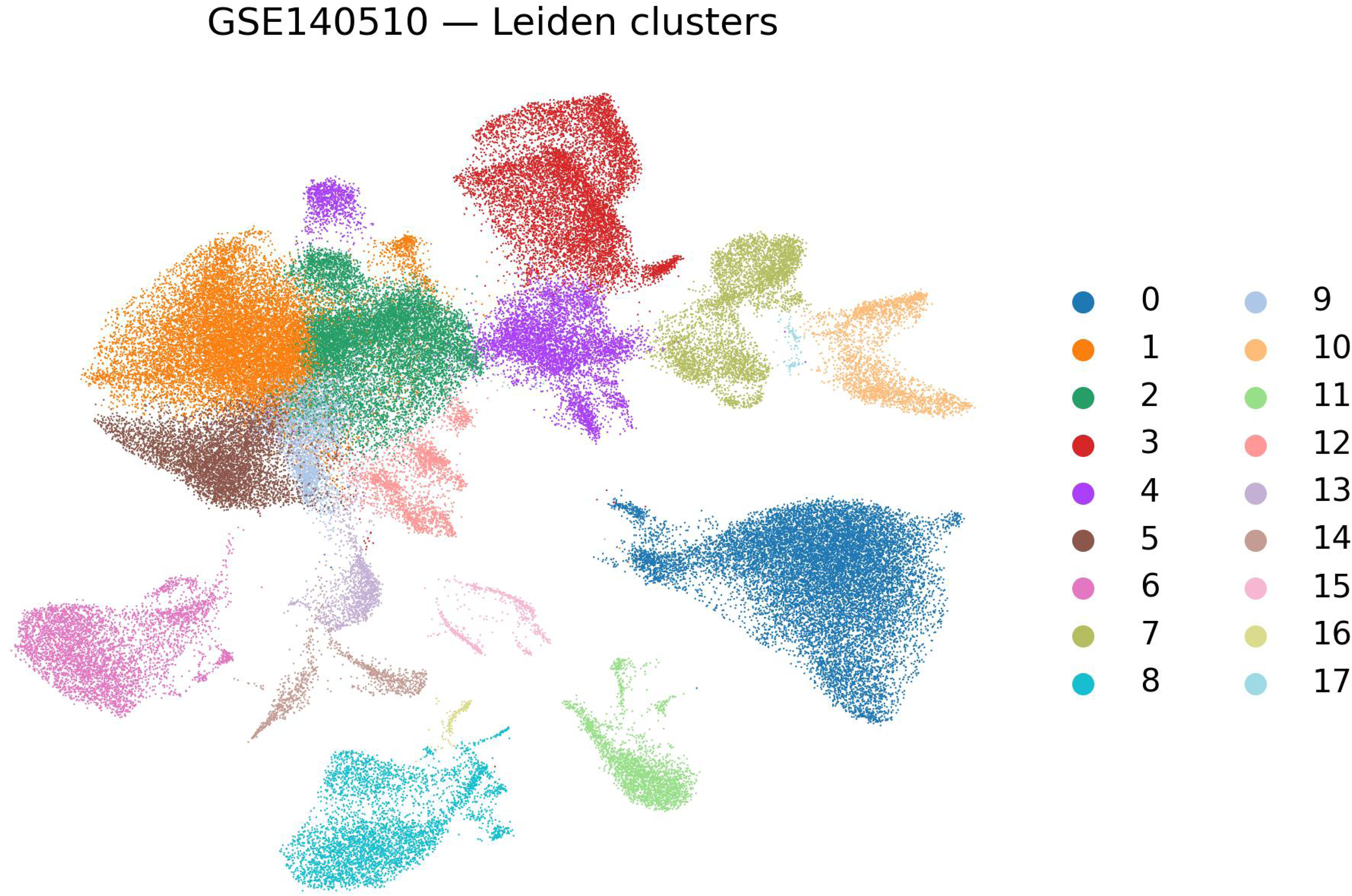

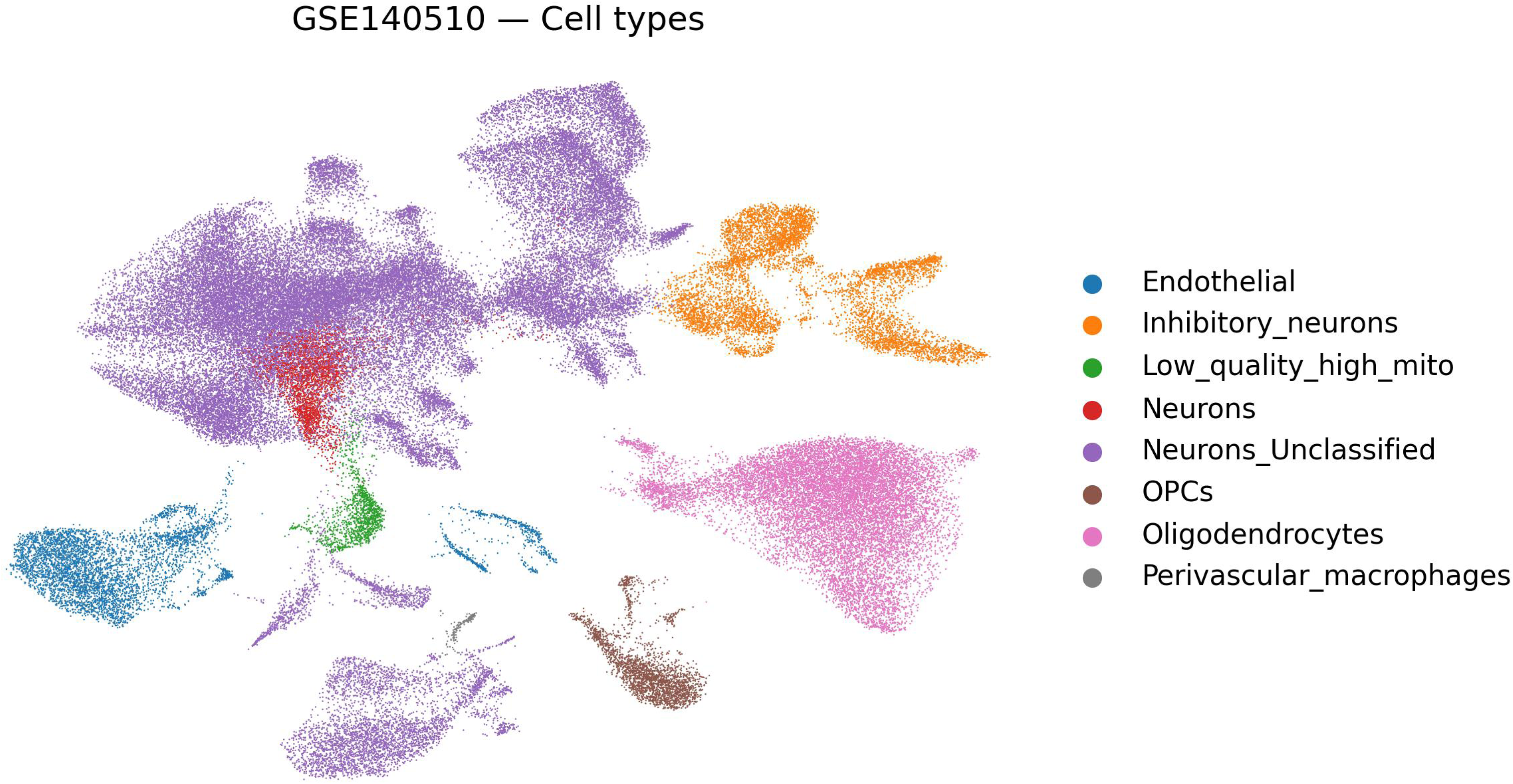

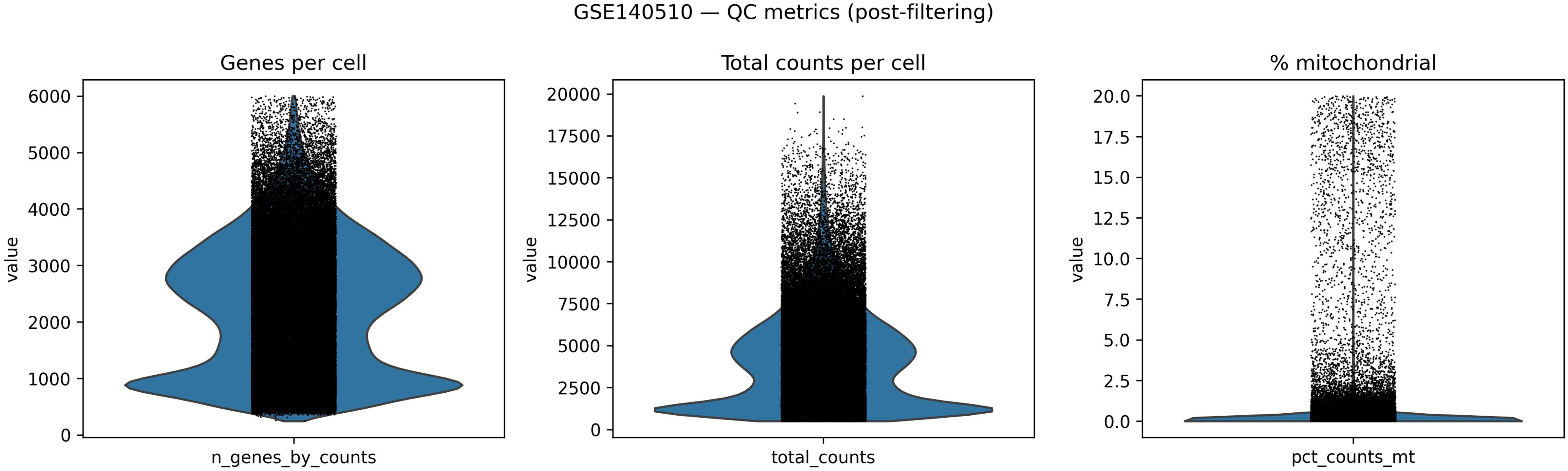

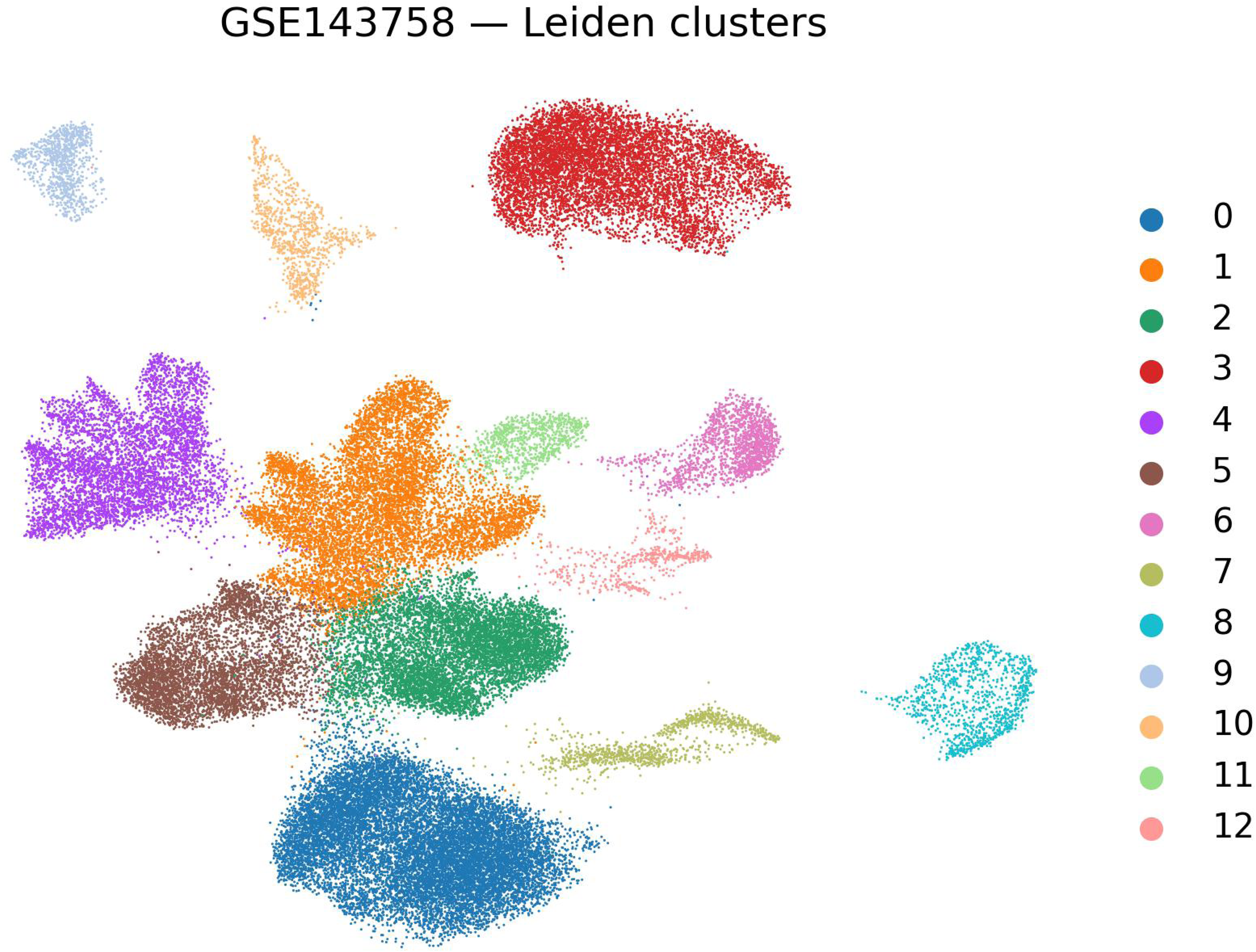

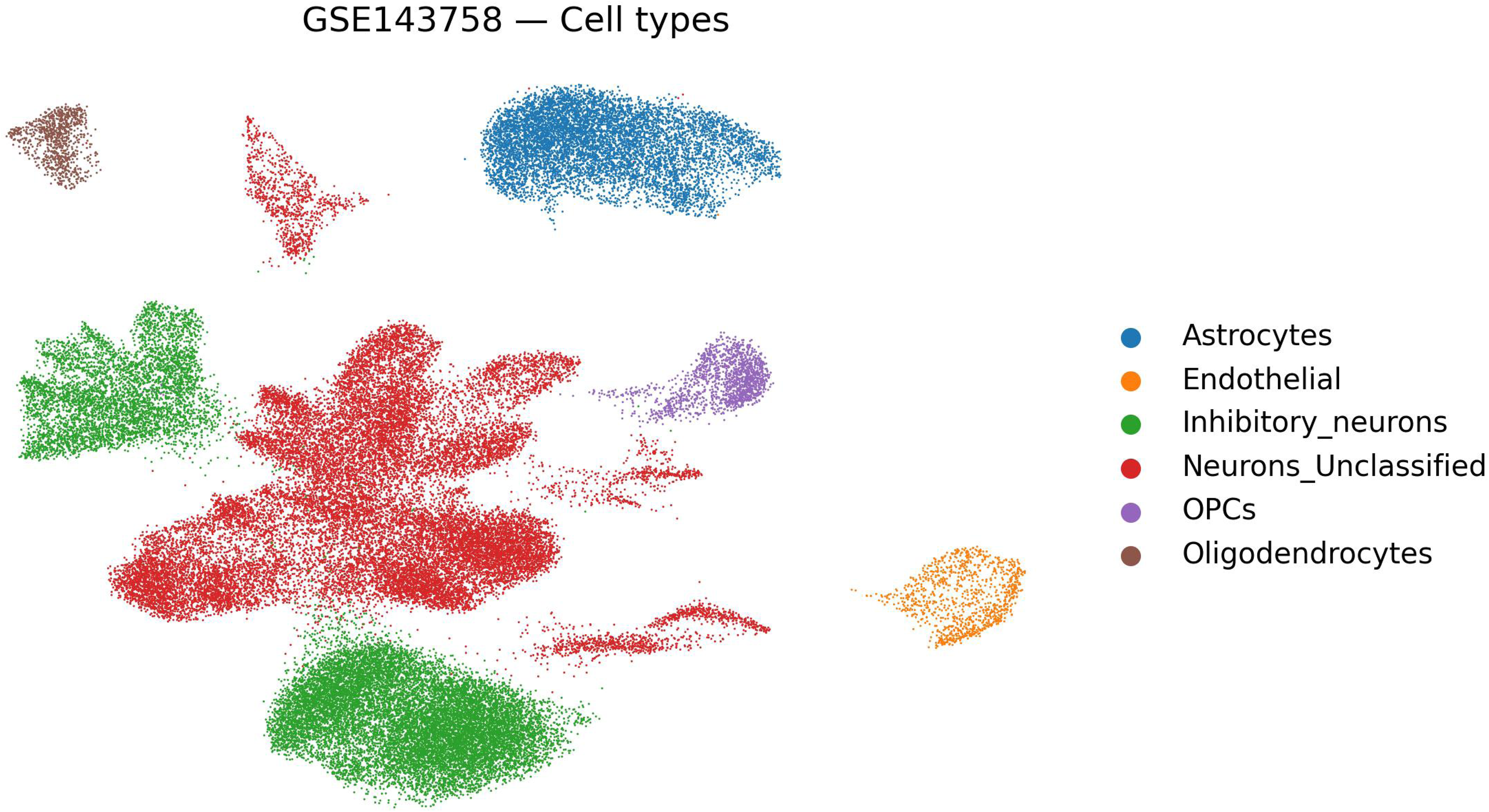

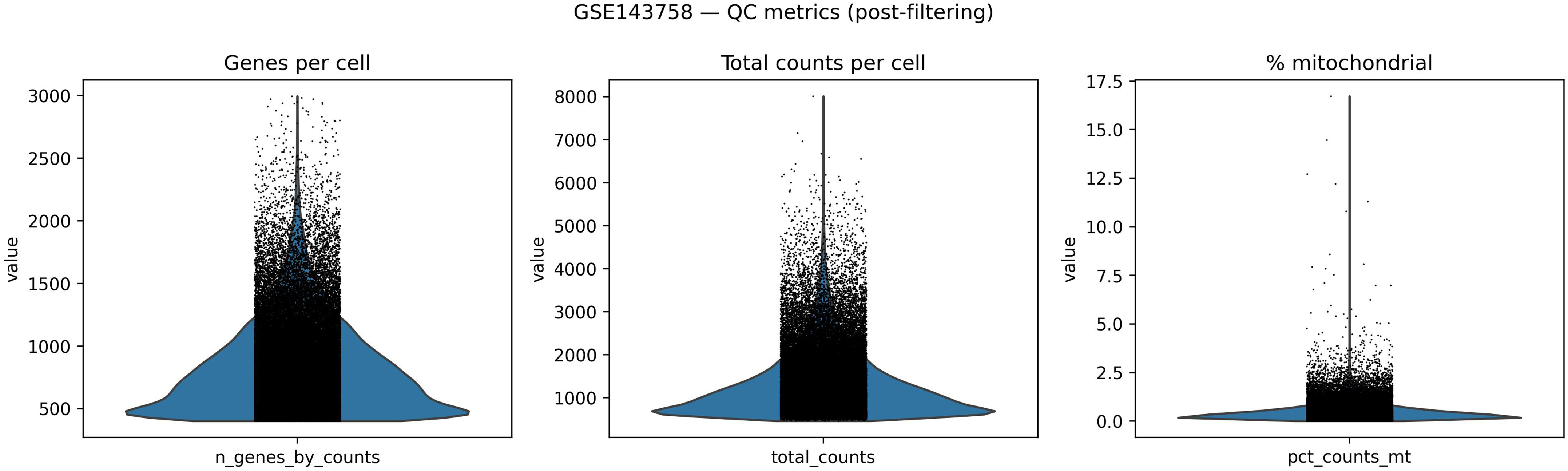

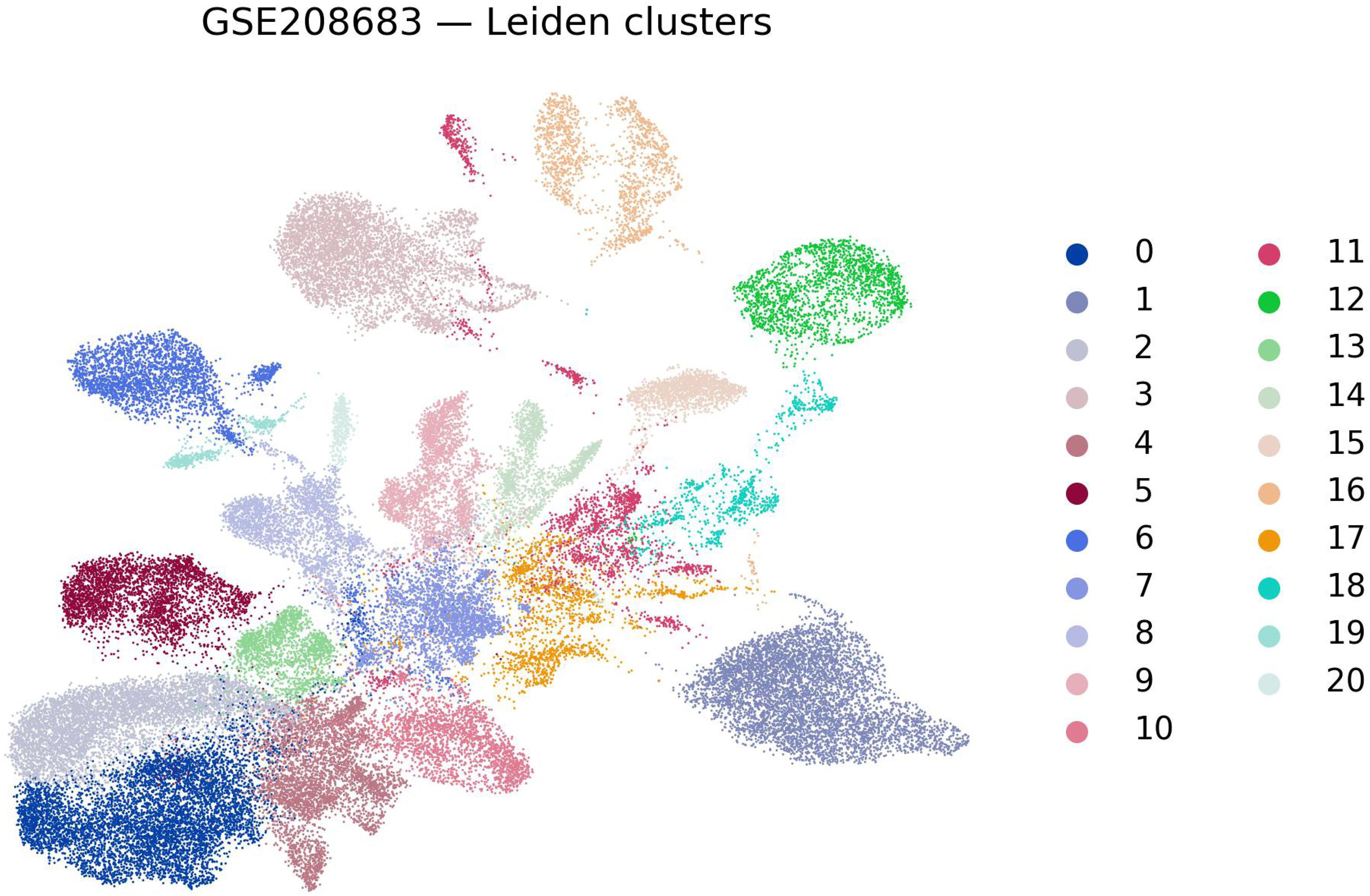

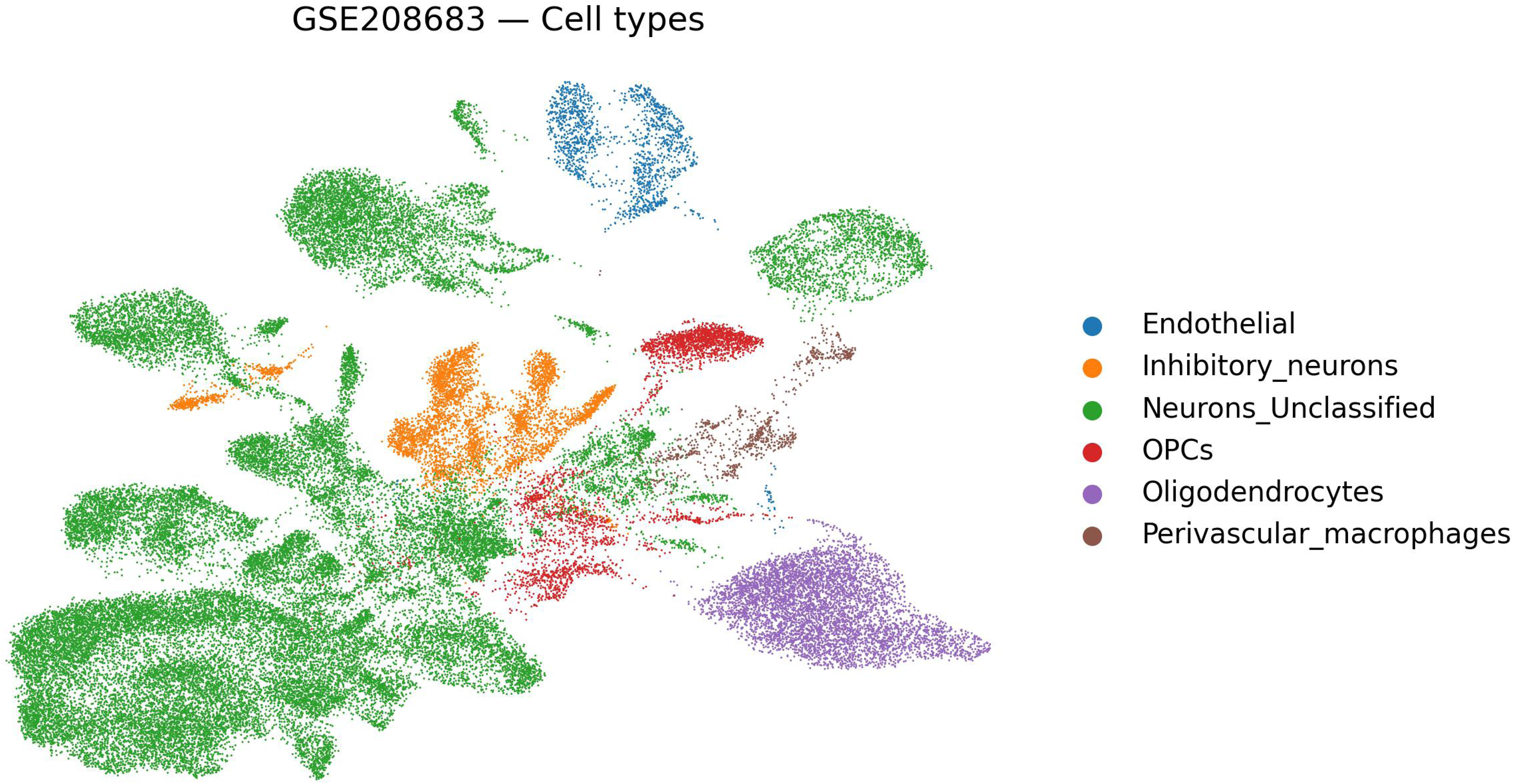

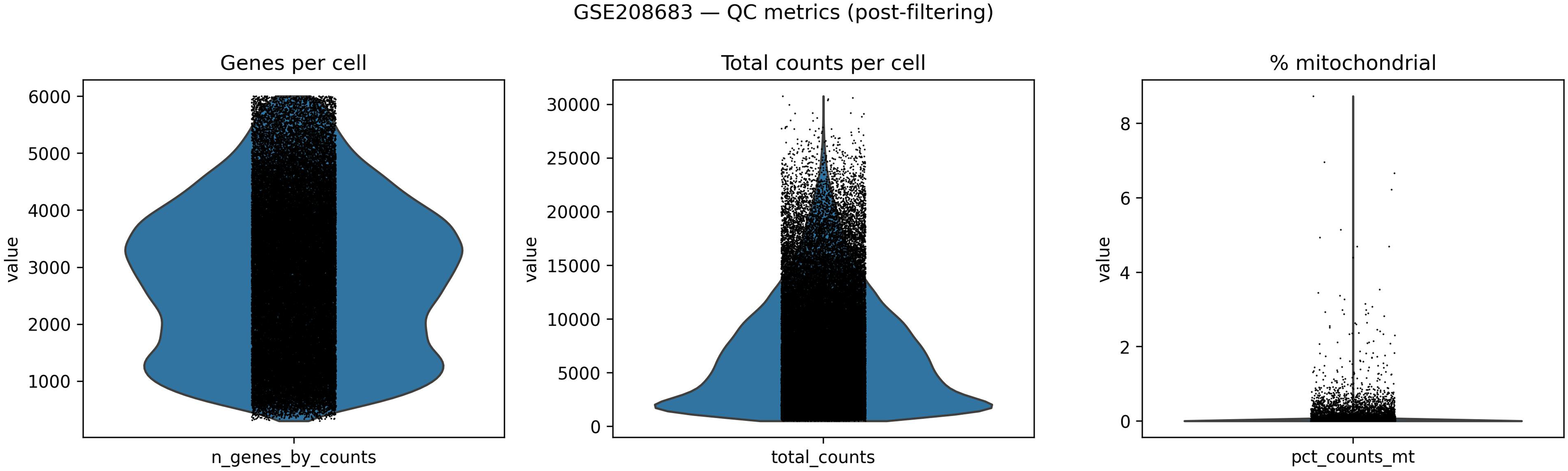

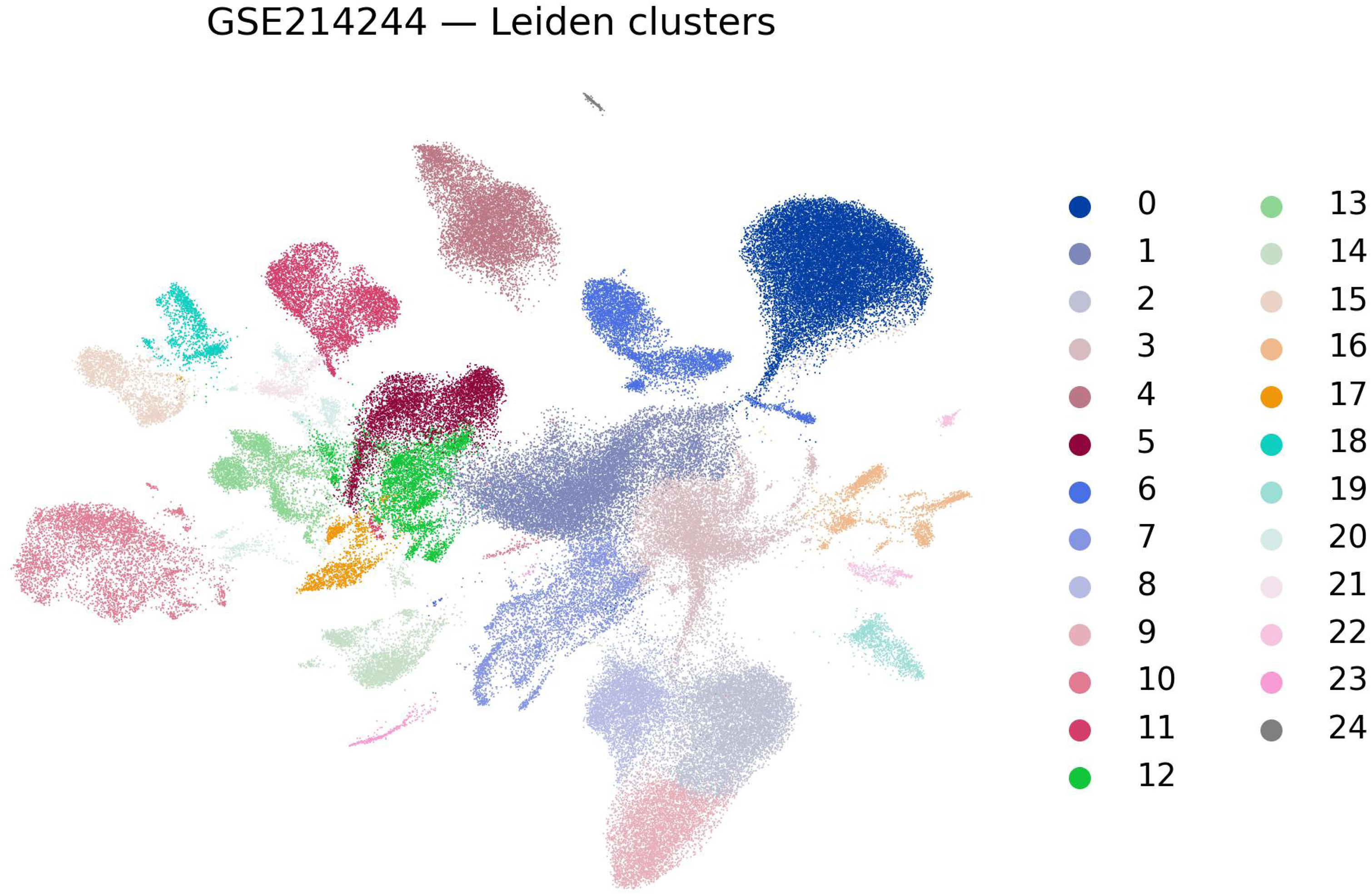

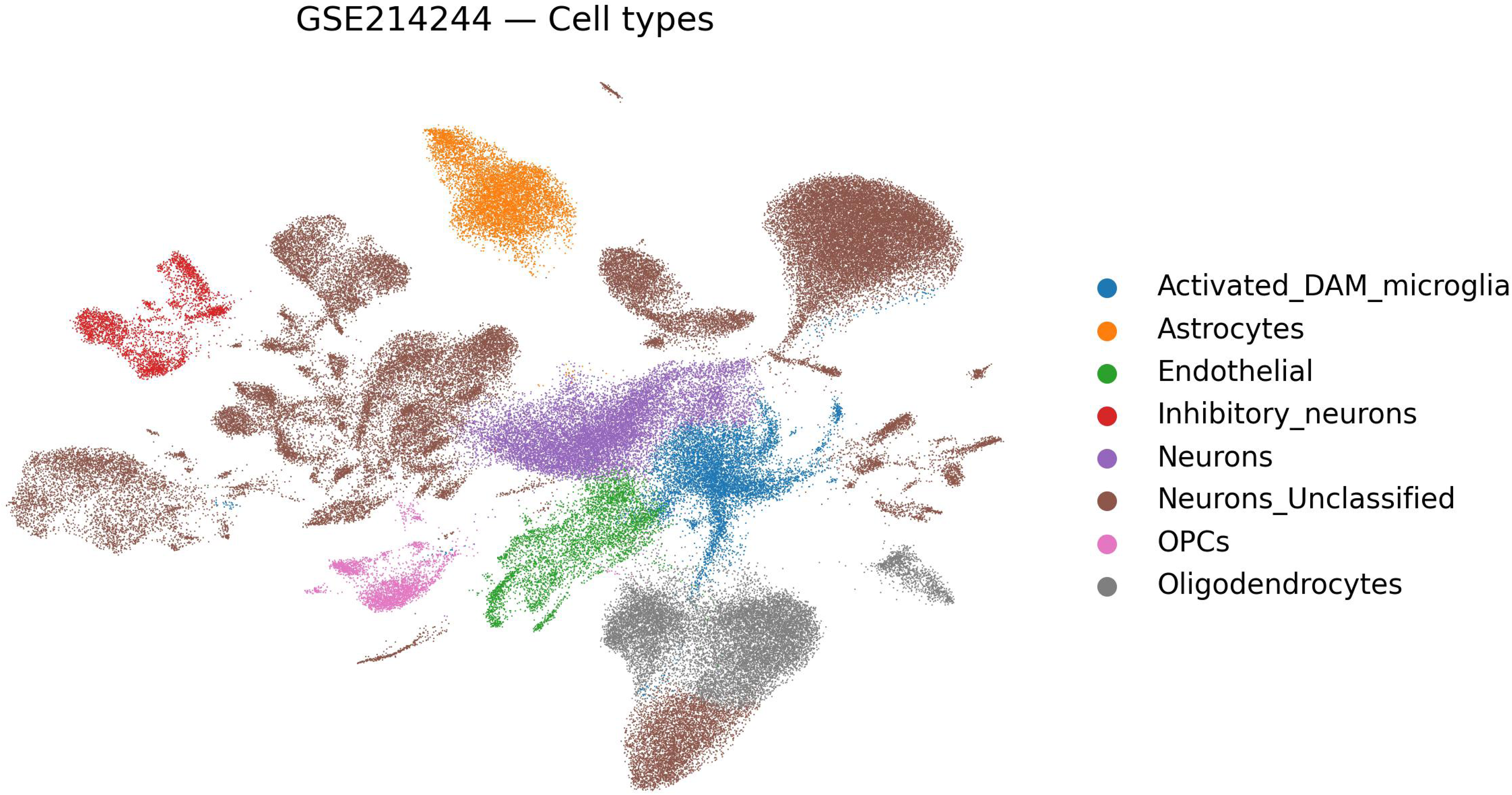

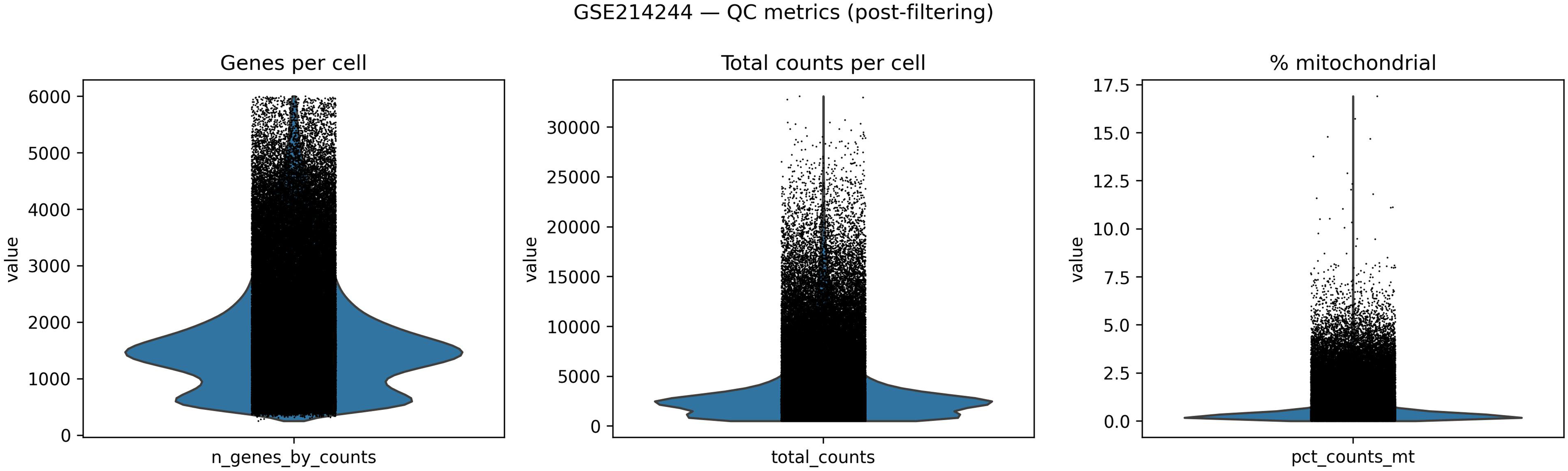

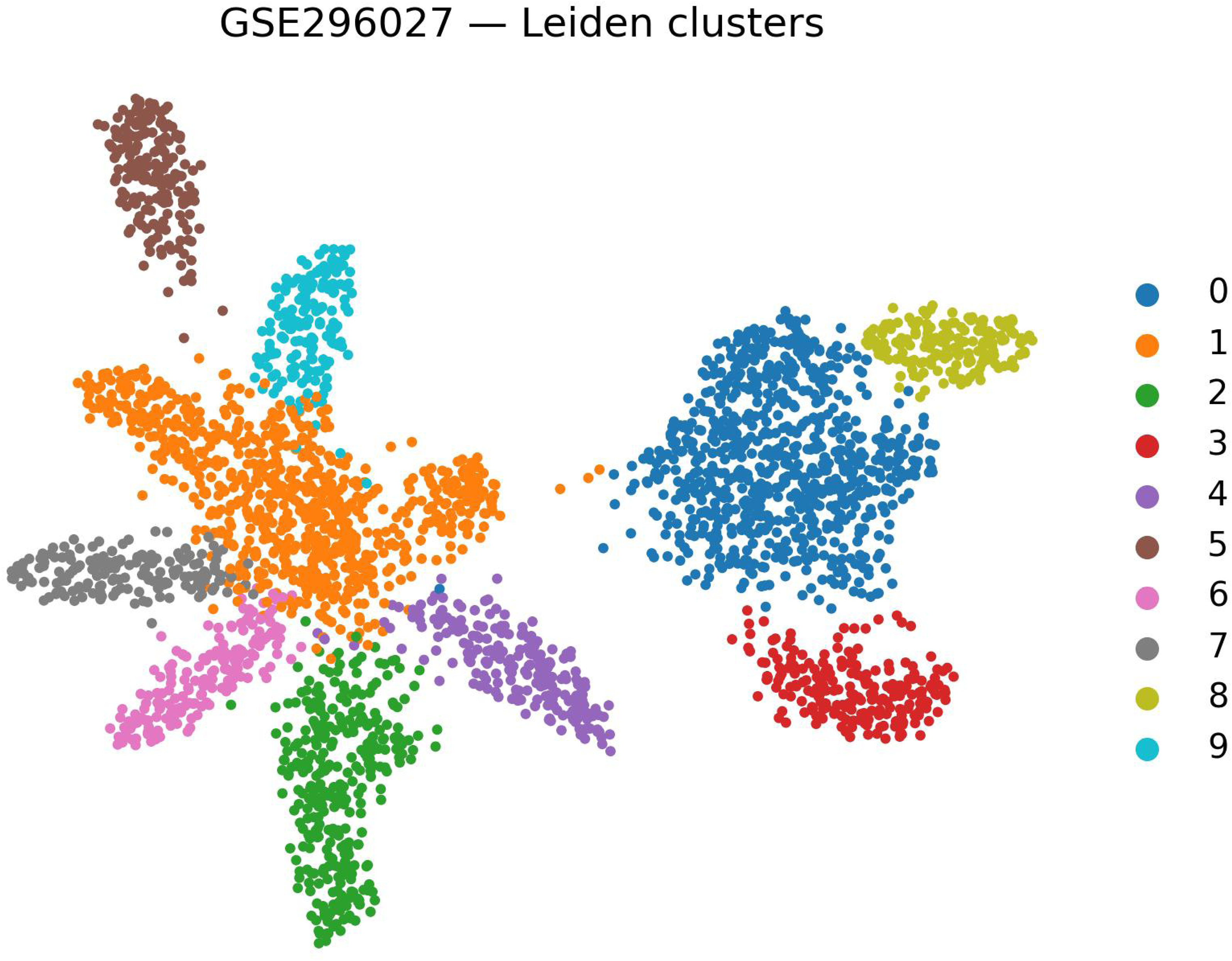

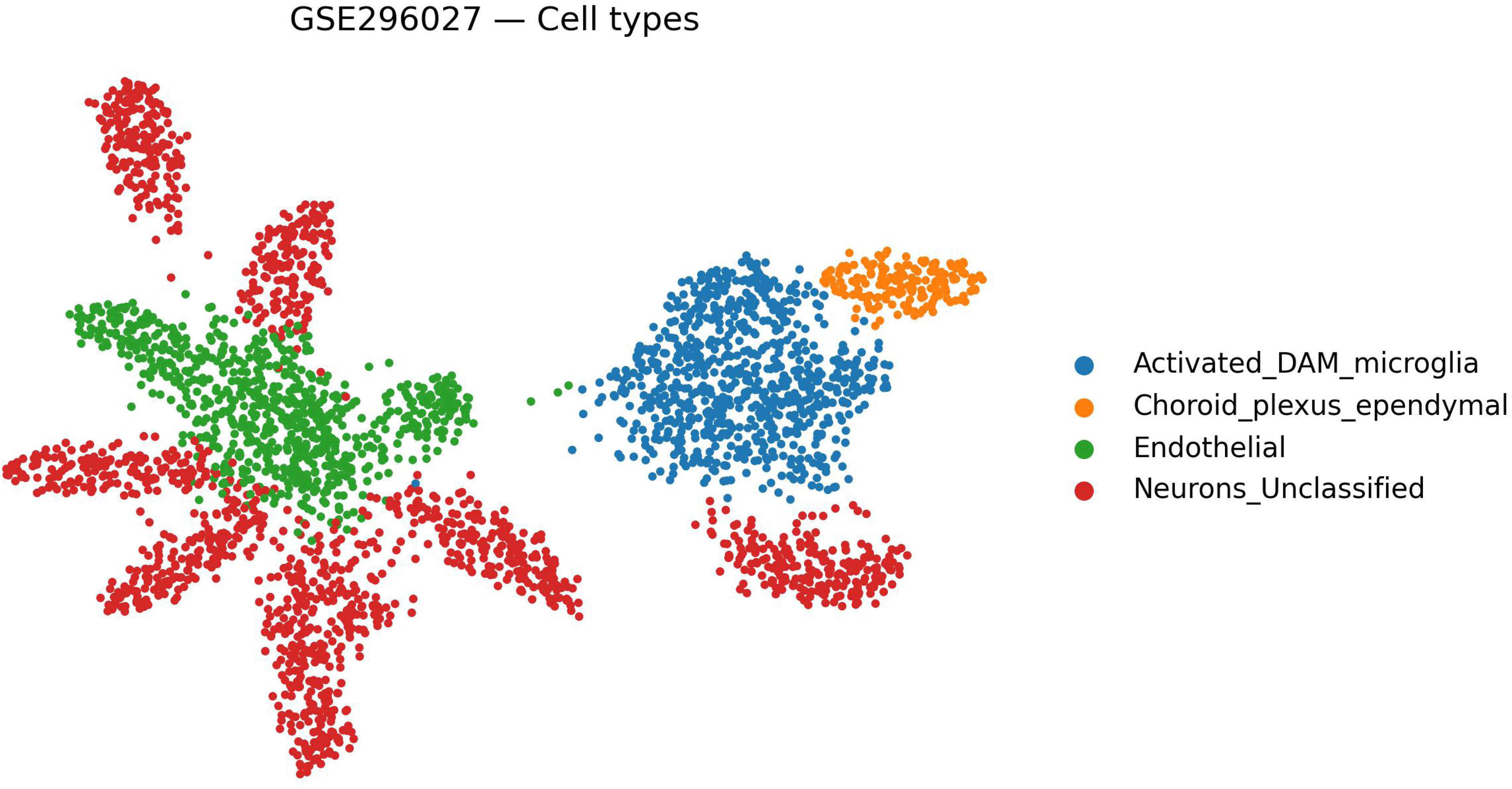

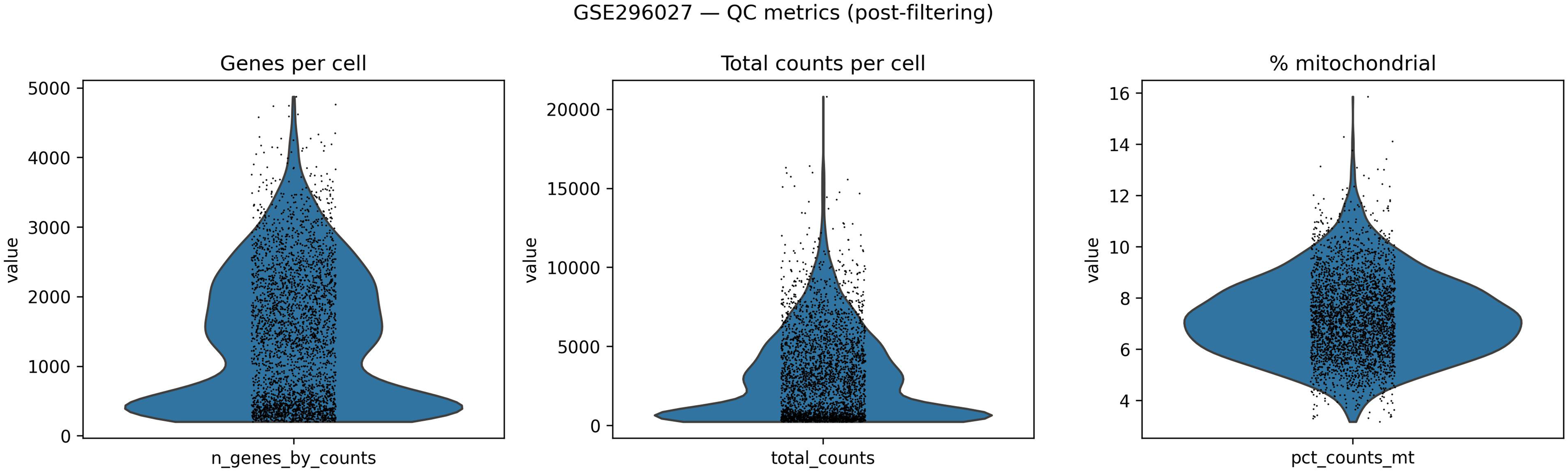

